# The developing hippocampus: Microstructural evolution through childhood and adolescence

**DOI:** 10.1101/2024.08.19.608590

**Authors:** Bradley G. Karat, Sila Genc, Erika P. Raven, Marco Palombo, Ali R. Khan, Derek K. Jones

## Abstract

The hippocampus is a structure in the medial temporal lobe which serves multiple cognitive functions. While important, the development of the hippocampus in the formative period of childhood and adolescence has not been extensively investigated, with most contemporary research focusing on macrostructural measures of volume. Thus, there has been little research on the development of the micron-scale structures (i.e., microstructure) of the hippocampus, which engender its cognitive functions. The current study examined age-related changes of hippocampal microstructure using diffusion MRI data acquired with an ultra-strong gradient (300 mT/m) MRI scanner in a sample of children and adolescents (N=88; 8-19 years). Surface-based hippocampal modelling was combined with established microstructural approaches, such as Diffusion Tensor Imaging (DTI) and Neurite Orientation Dispersion Density Imaging (NODDI), and a more advanced gray matter diffusion model Soma And Neurite Density Imaging (SANDI). No significant changes in macrostructural measures (volume, gyrification, and thickness) were found between 8-19 years, while significant changes in microstructure measures related to neurites (from NODDI and SANDI), soma (from SANDI), and mean diffusivity (from DTI) were found. In particular, there was a significant increase across age in neurite MR signal fraction and a significant decrease in extracellular MR signal fraction and mean diffusivity across the hippocampal subfields and long-axis. A significant negative correlation between age and MR apparent soma radius was found in the subiculum and CA1 throughout the anterior and body of the hippocampus. Further surface-based analyses uncovered variability in age-related microstructural changes between the subfields and long-axis, which may reflect ostensible developmental differences along these two axes. Finally, correlation of hippocampal surfaces representing age-related changes of microstructure with maps derived from histology allowed for postulation of the potential underlying microstructure that diffusion changes across age may be capturing. Overall, distinct neurite and soma developmental profiles in the human hippocampus during late childhood and adolescence are reported for the first time.

## 1. Introduction

The hippocampus is a widely studied yet enigmatic archicortical region that is typically parcellated into mesoscopic subfields which differ in both structure and function (Ding & Van Hoesen, 2015; Duvernoy et al., 2013). Part of its mystery arises from its relatively uncharacterized development. One hypothesis posits that the evolutionary development of the neocortex arose from a primordial hippocampus and amygdala, highlighting its importance in acquiring higher-order cognitive functions (Giaccio, 2006; Sanides, 1964; Sanides, 1970). However, little is known about how the hippocampus develops on the timescale of a human lifespan, particularly in late childhood (∼6-12 years) and adolescence (∼12-18 years). This characterization is critical to better understand the formation of human cognition, and the principal role the hippocampus has in functions like episodic and semantic memory, spatial navigation, emotion, behaviour, and more (Buzsáki & Moser, 2013; Squire, 1992; Strange et al., 2014; Sweatt, 2010).

While important, it is a challenging task to study the hippocampus during the early years of human development. Contemporary research has used non-invasive methods such as magnetic resonance imaging (MRI) to examine the hippocampus and its subfields across these early developmental stages (Callow et al., 2020; Giedd et al., 1996; Gogtay et al., 2006; Krogsrud et al., 2014; Langnes et al., 2020; Lee et al., 2014; Pfluger et al., 1999; Tamnes et al., 2018; Uematsu et al., 2012; Vinci-Booher et al., 2023; Wierenga et al., 2014). Most studies have focused on volumetric changes, where it has generally been shown that hippocampal volume increases across childhood and adolescence, likely capturing an expansion of cognitive capacity (Krogsrud et al., 2014; Langnes et al., 2020; Lee et al., 2014; Pfluger et al., 1999; Tamnes et al., 2018; Wierenga et al., 2014). However, there have been conflicting results, with studies finding variable patterns of age-related hippocampal volume changes (Giedd et al., 1996; Lee et al., 2014; Tamnes et al., 2018; Wierenga et al., 2014). While volume does appear to be sensitive to developmental changes, it is a coarse measure which is unspecific towards the intrahippocampal gray matter. This includes components such as glial cells, neurites, soma and other micron-scale structures (collectively termed microstructure) that are responsible for the computations which engender hippocampal function and are of critical importance in both health and disease.

The development of gray matter microstructure is generally characterized by rapid growth of dendrites, axons, and a proliferation of synaptic connections in early childhood, which are pruned during later childhood years (Goldman-Rakic, 1987). Diffusion MRI (dMRI) is a technique which sensitizes the MRI signal to the micron-scale movement of water (Le Bihan, 1995) which can be leveraged to study developmental patterns of microstructure (Lebel et al., 2008). Previous studies using Diffusion Tensor Imaging (DTI; Basser et al., 1994) to capture hippocampal microstructural development have generally found a negative correlation between mean diffusivity and age in early childhood, while age-related changes in fractional anisotropy have been more variable (Callow et al., 2020; Langnes et al., 2020; Vinci-Booher et al., 2023). However, these findings are not specific to any particular microstructural property, and could be a result of changes to axon or dendrite density, myelination, soma related changes, or other micron level alterations (Jensen & Helpern, 2010; Jelescu & Budde, 2017; Karat et al., 2024). Recent advances in MRI hardware including stronger gradients (Jones et al., 2018; McNab et al., 2013; Setsompop et al., 2013) and new modelling approaches (Palombo et al., 2020) appear promising to disentangle apparent soma and neurite contributions to the dMRI signal in vivo.

In this work we examined age-related changes of hippocampal microstructure using dMRI data acquired using an ultra-strong gradient (300 mT/m) MRI scanner in a sample of children and adolescents (aged 8-19 years; Genc et al., 2020; Genc et al., 2024). Using HippUnfold, a hippocampal surface-based approach, we investigated age and sex-related changes of macro- and microstructure across the subfields and long-axis (DeKraker et al., 2022). In particular, the Soma And Neurite Density Imaging (SANDI; Palombo et al., 2020) model was used to derive measures related to both the soma and neurite. The Neurite Orientation Dispersion and Density Imaging model (NODDI; Zhang et al., 2012) and DTI (Basser et al., 1994) were also used to compare microstructure trajectories across age. Utilizing the salient orientation information from dMRI, we also determined if there were shifts in diffusion orientation across age which may be related to the development of the complex but organized intrahippocampal circuitry. Finally, we derived surface maps which capture age effects for all macro- and microstructural measures. We then correlated these maps with metrics derived from histology and high-resolution MRI to postulate what the age-related changes might be capturing in terms of known microstructure. Overall, we report distinct neurite and soma developmental profiles in the human hippocampus during late childhood/adolescence for the first time. This forms a crucial baseline for understanding the structural alterations underlying the progressive formation of human cognition and developmental disorders, as well as open new avenues for corroborating in vivo diffusion with histology.

## 2. Results

### 2.1 Age and sex-related changes in subfield volume & macrostructure

Previous research has typically analyzed the volume of the hippocampus across development (Krogsrud et al., 2014; Langnes et al., 2020; Lee et al., 2014; Uematsu et al., 2012). Figure 1 depicts the correlation between age and subfield averaged macrostructural measures of volume, gyrification, thickness. No significant interaction was found between age and hemisphere for volume (F(1,864)=0.001, p=0.974), gyrification (F(1,864)=0.104, p=0.747), or thickness (F(1,864)=0.554, p=0.457), suggesting that the hemispheres display similar age-related changes in subfield macrostructure (supplementary figure S1). Thus hemisphere data (i.e., between left and right hippocampus) was averaged within participants.

**Figure 1.**
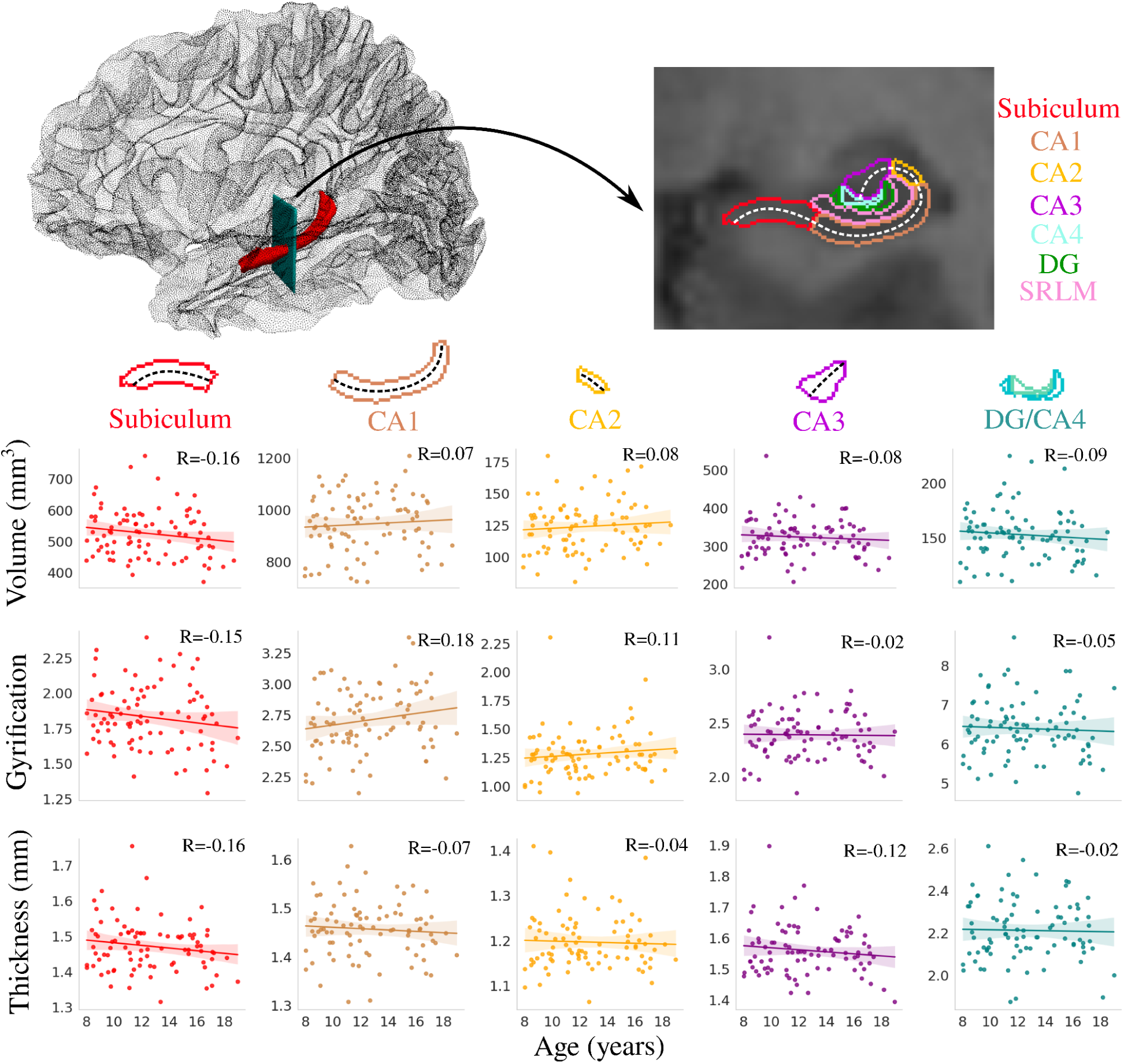
Correlation between age and subfield averaged macrostructural measures of volume, gyrification, and thickness. The top left figure depicts the 3D location of the hippocampus (shown in red), and the arrow represents the position of the coronal slice shown on the top right figure. Colours represent hippocampal subfields and relationships are quantified using Pearson’s correlation coefficient (R). The dashed lines approximately represent the midthickness surface which gyrification and thickness were calculated and then averaged on (note that the surface excludes the SRLM). As well, the DG and CA4 were averaged together. CA - cornu ammonis; DG - dentate gyrus; SRLM - stratum radiatum lacunosum moleculare.

No significant correlations were found between age and any subfield averaged macrostructural measure (figure 1). Similar results were found for anterior-posterior averaged macrostructure (supplementary figure S2). As well, no significant correlation between age and subfield volume was found using FreeSurfer (supplementary figure S3), corroborating the result seen in figure 1.

Figure 2 depicts the correlation between age and subfield averaged macrostructural measures of volume, gyrification, thickness stratified by sex. Interestingly, it appears that males generally have positive correlations between age and subfield volume, gyrification, and thickness, while females showed little changes across age. The interaction between age and sex was significant after false-discovery rate (FDR) correction for volume (F(1,424)=6.03, p-adjusted=0.040) and thickness (F(1,424)=4.966, p-adjusted=0.040), while not significant for gyrification (F(1,424)=1.573, p-adjusted=0.211).

**Figure 2.**
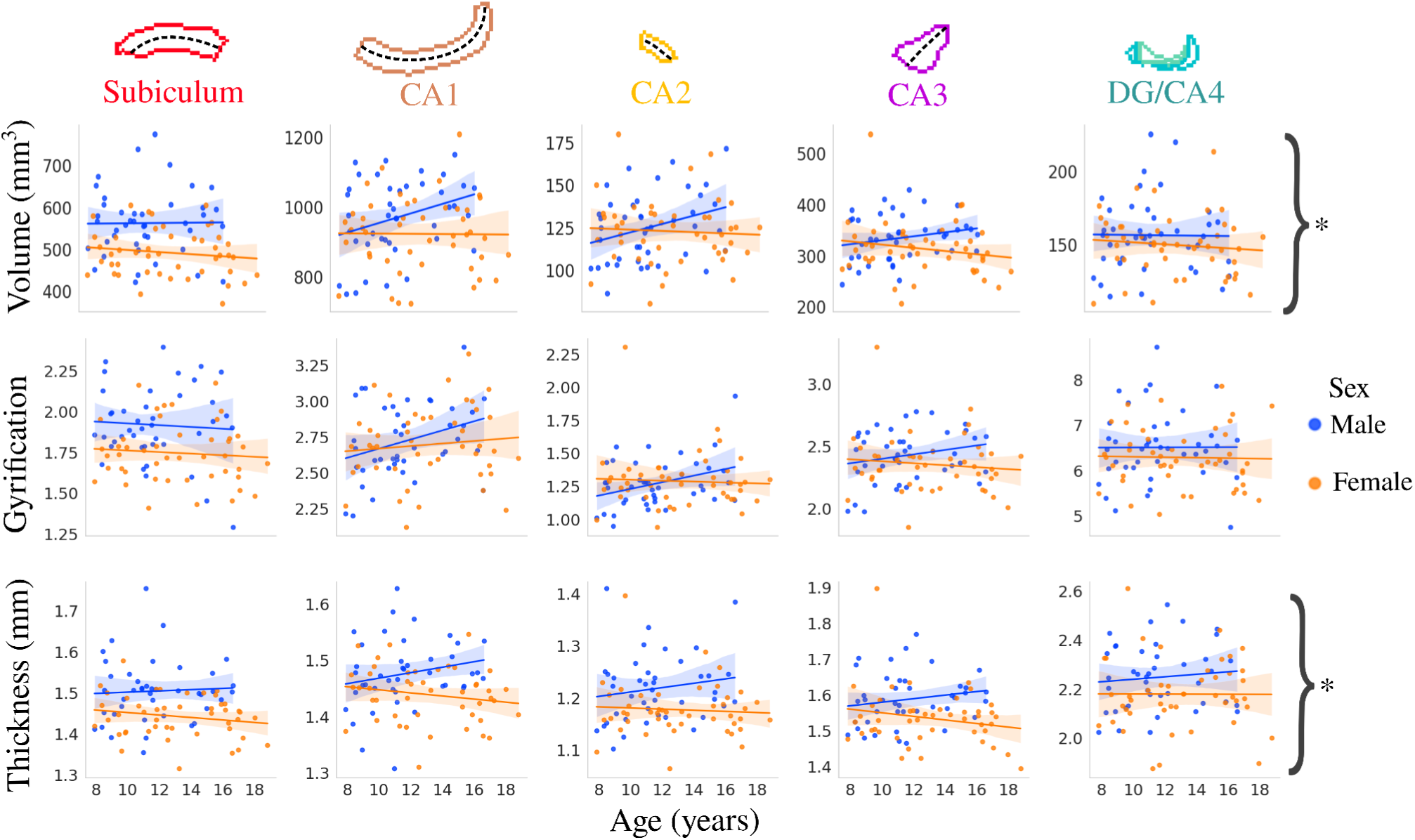
Relationship between age and macrostructure by hippocampal subfield and stratified by sex. Asterisks represent metrics with a significant interaction between age and sex after FDR correction.

### 2.2 Age and sex-related changes in subfield microstructure

Supplementary figure S4 depicts the relationship between age and subfield-averaged partial volume maps of CSF, GM, and WM (see *section 5.2)*. No significant correlations were found between age and the CSF partial volume measure. In CA1, CA2, CA3, and the DG/CA4, the GM tissue probability is generally between 0.8-1 while the CSF probability is between 0-0.07, suggesting that the microstructure measures sampled on the midthickness surface are mostly within the GM. The subiculum has a higher WM tissue probability (0.2-0.4), which is expected given the presence of the highly myelinated perforant path.

Figure 3 depicts the correlation between subfield averaged microstructural measures and age using SANDI (neurite, soma, and extracellular MR signal fractions as well as MR apparent soma radius), NODDI (orientation dispersion index), and DTI (mean diffusivity) metrics. No significant interaction was found between age and hemisphere for fneurite_SANDI_ (F(1,864)=0.014, p=0.906), fsoma (F(1,864)=0.001, p=0.971), fextracellular (F(1,864)=0.014, p=0.907), Rsoma (F(1,864)=0.576, p=0.448), ODI (F(1,864)=0.089, p=0.766), and MD (F(1,864)=1.223, p=0.269), suggesting that the hemispheres display similar age-related changes in subfield microstructure (supplementary figure S5). Thus hemisphere data was averaged within participants.

**Figure 3.**
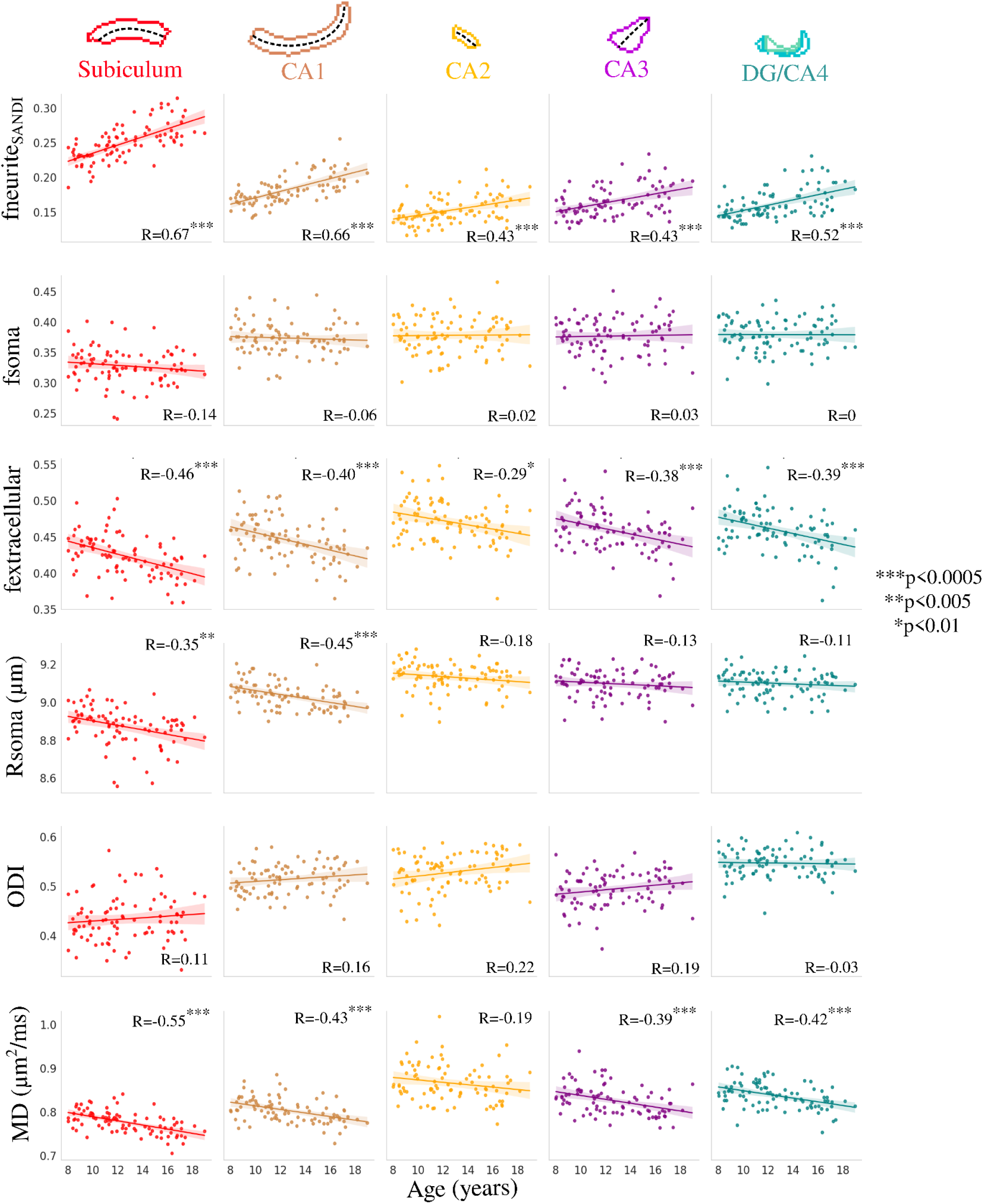
Correlation between age and subfield averaged microstructural measures of neurite (fneurite_SANDI_), soma (fsoma), and extracellular (fextracellular) MR signal fractions, soma radius (Rsoma), orientation dispersion index (ODI) and mean diffusivity (MD). Colours represent hippocampal subfields and relationships are quantified using Pearson’s correlation coefficient (R). The dashed lines approximately represent the midthickness surface which the metrics were sampled and then averaged on. CA - cornu ammonis; DG - dentate gyrus.

Many significant correlations were found between age and subfield-averaged microstructural measures (figure 3). In particular, fneurite_SANDI_ was significantly positively correlated with age across all subfields while fextracellular and MD were significantly negatively correlated with age across all subfields (with the exception of CA2; figure 3). Rsoma was significantly negatively correlated with age only in the subiculum and CA1. ODI and fsoma were not significantly correlated with age in any subfield. Interestingly, the age-related slopes seen in figure 3 were significantly different across the subfields for fneurite_SANDI_ (F(4,424)=4.354, p-adjusted=0.011) and Rsoma (F(4,424)=3.086, p-adjusted=0.047), suggesting that the subfields may have unique patterns of neurite and soma development. However, the slopes across age were not significantly different across the subfields for fsoma (F(4,424)=0.511, p-adjusted=0.894), fextracellular (F(4,424)=0.274, p-adjusted=0.894), ODI (F(4,424)=0.838, p-adjusted=0.894), and MD (F(4,424)=0.281, p-adjusted=0.894).

Figure 4 depicts the correlation between subfield averaged microstructural measures and age stratified by sex. The interaction between age and sex was significant after false-discovery rate correction for fneurite_SANDI_ (F(1,424)=43.39, p-adjusted=8×10^-10^), fextracellular (F(1,424)=4.509, p-adjusted=0.04), Rsoma (F(1,424)=16.819, p-adjusted=1.47×10^-04^), ODI (F(1,424)=10.392, p-adjusted=0.002), and MD (F(1,424)=14.247, p-adjusted=3.66×10^-04^). The age-by-sex interaction was not significant for fsoma (F(1,424)=2.673, p-adjusted=0.103).

**Figure 4.**
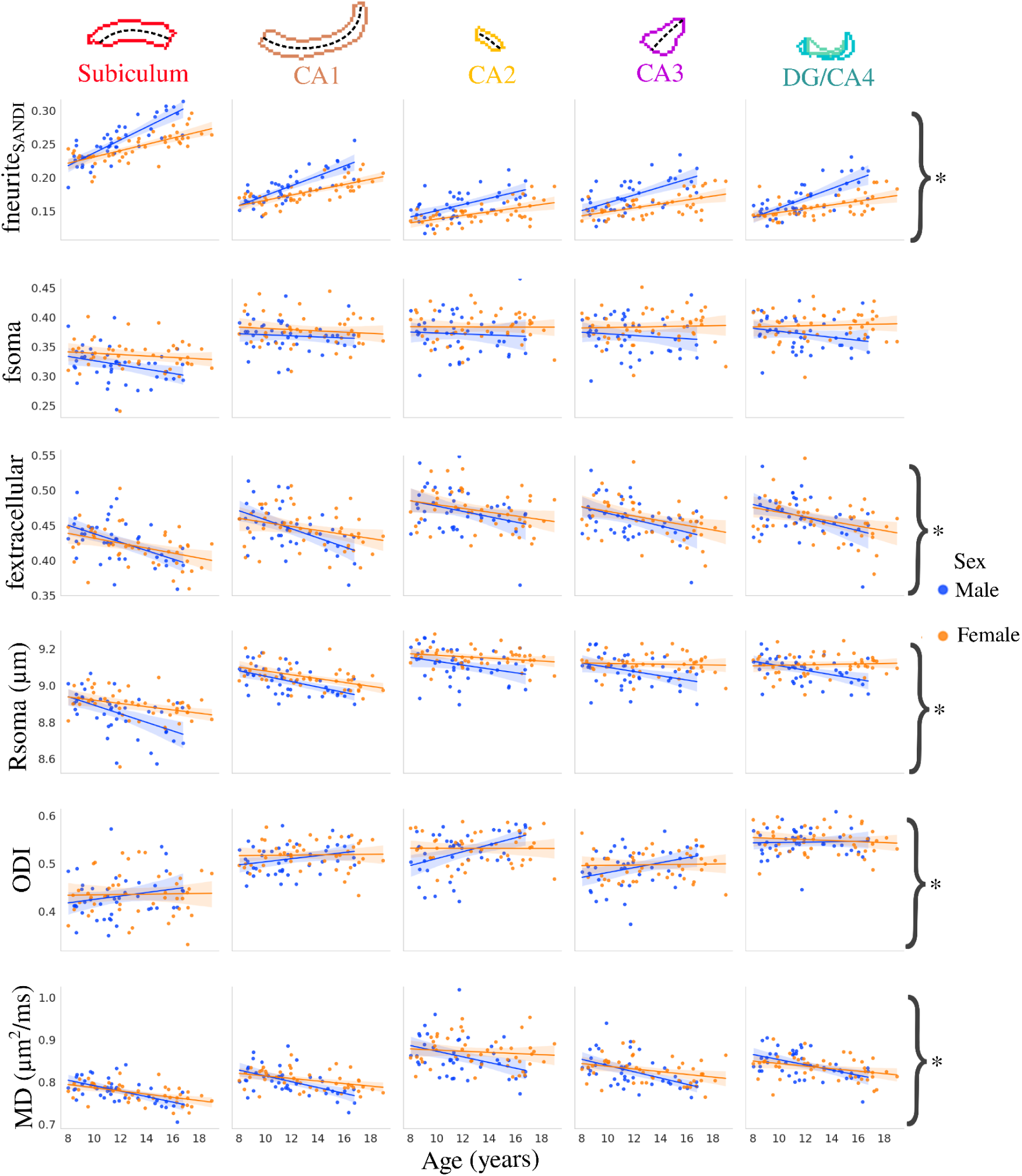
Relationship between age and microstructure by hippocampal subfield and stratified by sex. Asterisks represent metrics with a significant interaction between age and sex after FDR correction.

### 2.3 Age-related changes in long-axis microstructure

Figure 5 depicts the correlation between long-axis averaged microstructural measures and age using the same metrics as *section 2.2*. fneurite_SANDI_ was significantly positively correlated with age within each long-axis parcellation, while fextracellular and MD were significantly negatively correlated with age (figure 5). Rsoma was significantly negatively correlated with age in the lateral anterior and body portions of the hippocampus, while ODI was significantly positively correlated with age only in the posterior portion of the hippocampal body. Unlike within the subfields, the age-related long-axis slopes were not significantly different across the parcellations for fneurite_SANDI_ (F(4,424)=0.323, p=0.862), fsoma (F(4,424)=0.292, p=0.883), fextracellular (F(4,424)=0.373, p=0.828), Rsoma (F(4,424)=0.804, p=0.523), ODI (F(4,424)=2.054, p=0.086), and MD (F(4,424)=1.126, p=0.344).

**Figure 5.**
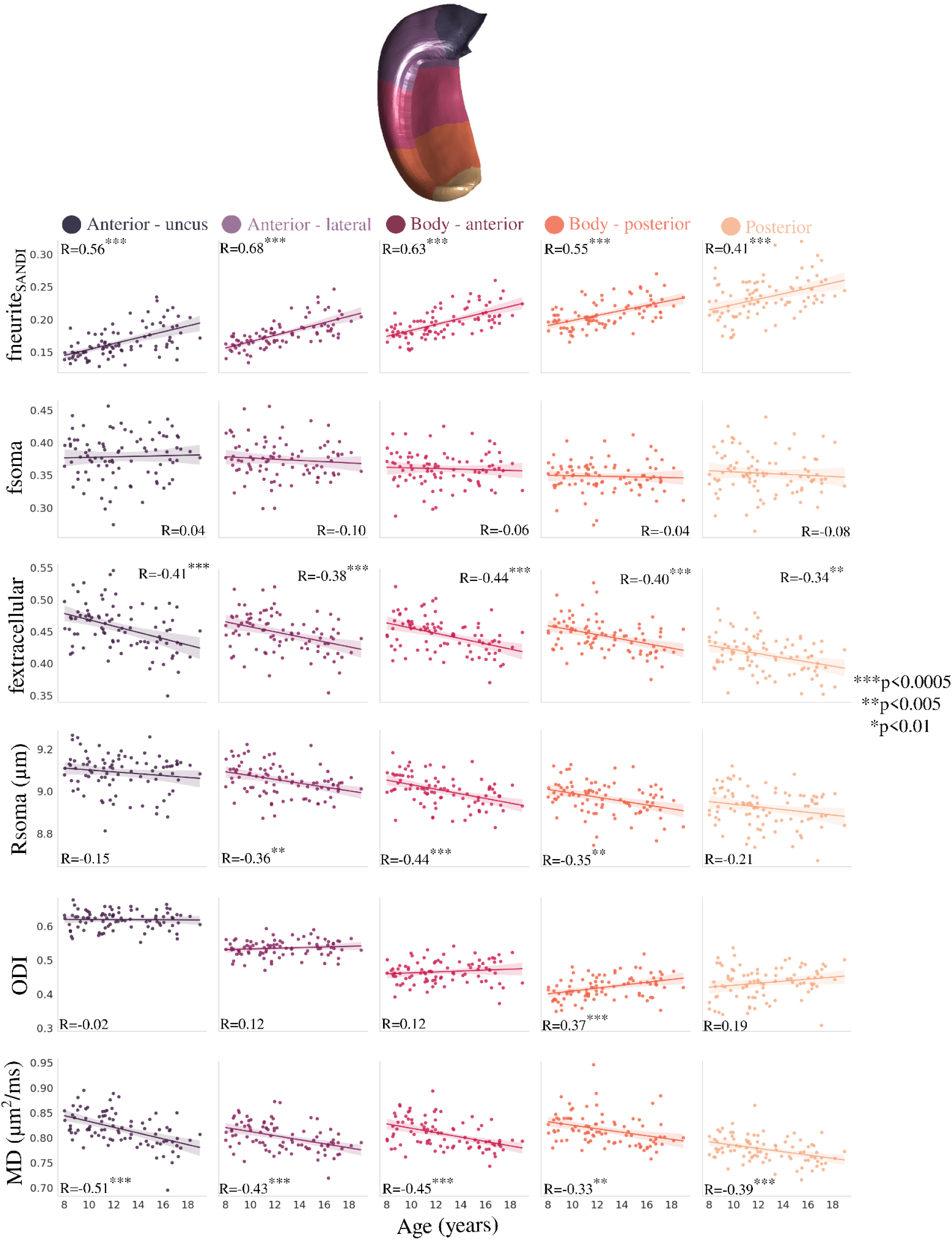
Correlation between age and anterior-posterior averaged microstructural measures. Colours represent hippocampal long-axis parcellations (shown on a midthickness surface at the top) and relationships are quantified using Pearson’s correlation coefficient (R).

### 2.4 Diffusion orientation changes across age

Using the first peak of the fiber orientation distribution function (see *section 5.4),* we quantified how diffusion orientations vary across age. Figure 6 depicts the correlation between age and subfield-averaged measures of long-axis, tangential, and radial oriented diffusion. Long-axis oriented diffusion was significantly positively correlated with age in CA1. The age-related slopes were not significantly different across the subfields for the long-axis orientations (F(4,424)=1.6, p=0.173), tangential orientations (F(4,424)=1.125, p=0.344), and radial orientations (F(4,424)=0.767, p=0.547).

**Figure 6.**
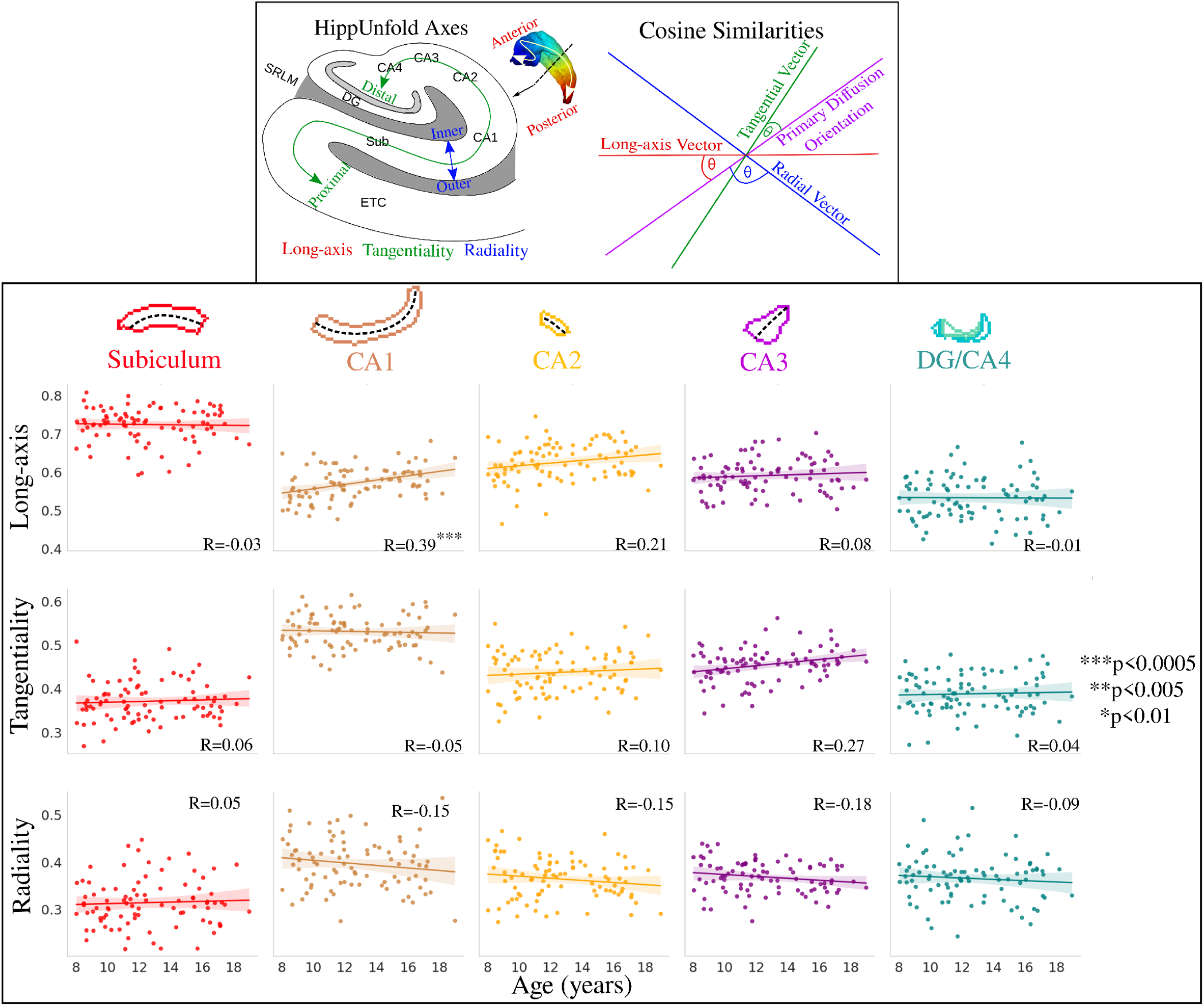
Relationship between age and diffusion orientations (cosine similarities) across the hippocampal subfields and quantified with pearson’s R. Top box (adapted from Karat et al., 2023) depicts the calculation of the cosine similarity, where the *HippUnfold* (DeKraker et al., 2023) axes are used to generate long-axis, tangential, and radial vectors which are then compared with the primary diffusion orientation (peak 1 from the fiber orientation distribution function).

Figure 7 depicts the same diffusion orientation metrics averaged across the long-axis parcellation. Long-axis oriented diffusion was significantly positively correlated with age in the lateral anterior region and the anterior hippocampal body. Radial oriented diffusion displayed a significant negative correlation with age in the lateral anterior region and a significant positive correlation with age in the posterior hippocampal body. This suggests that the diffusion orientation changes across age are occurring more in the anterior and body of the hippocampus. Interestingly, unlike across the subfields, the age-related slopes in figure 7 were significantly different across the long-axis parcellations for both long-axis (F(4,424)=3.190, p-adjusted=0.020) and radial (F(4,424)=5.341, p-adjusted=0.001) oriented diffusion.

**Figure 7.**
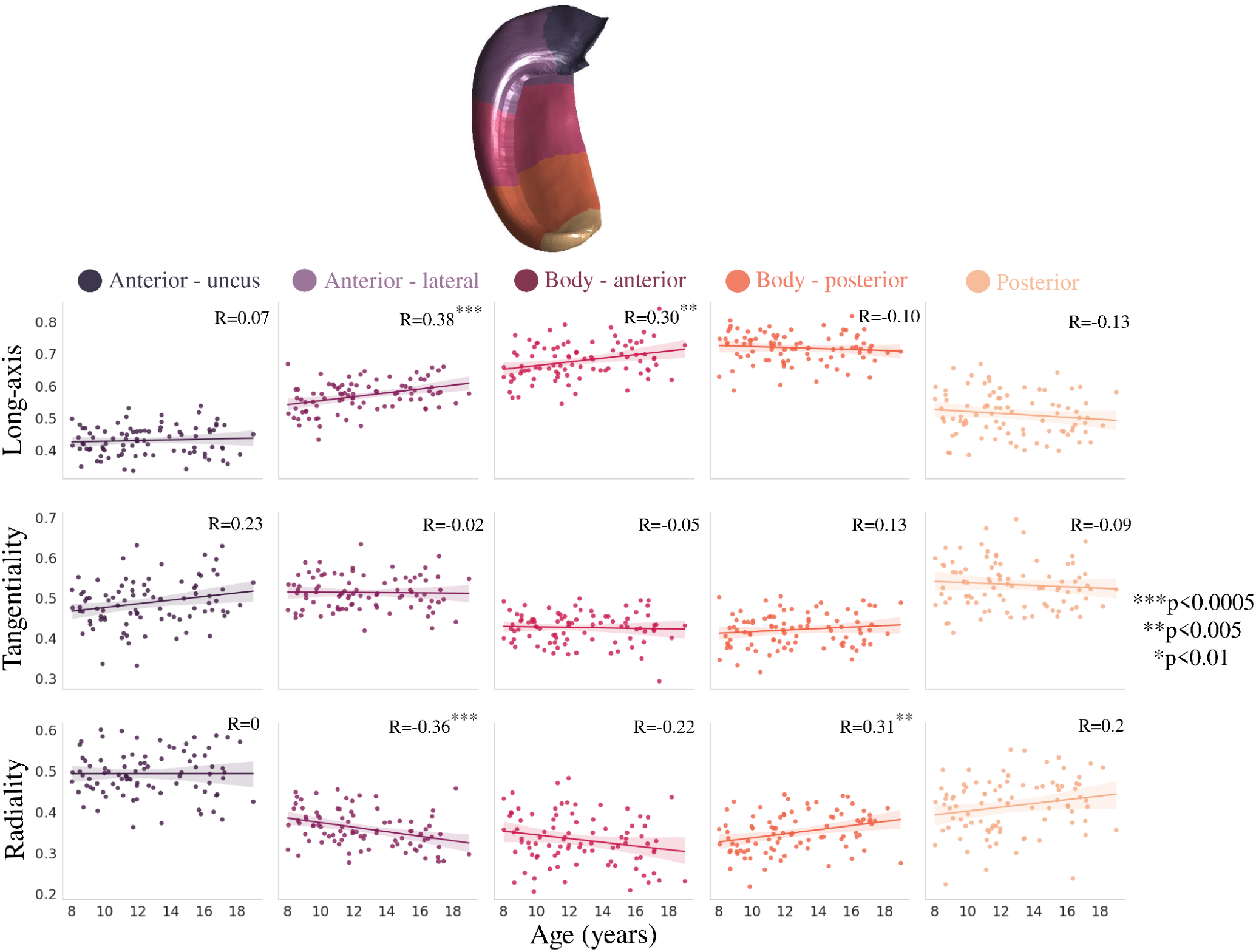
Relationship between age and diffusion orientations (cosine similarities) across the hippocampal long-axis and quantified with Pearson’s R.

### 2.5 Vertex-wise relation between macro- and microstructure across age and the hippocampal proximal-distal and long-axis

Surface t-statistic maps which capture the spatial changes in macro- and microstructural development were generated (figure 8B; see *section 5.5).* These age contrast maps were then correlated with contrived positional AP and PD gradients (figure 8A; DeKraker et al., 2024). The correlation then describes along which hippocampal axis are any age-related macro- or microstructural changes occurring (figure 8C). For example, the age contrast map of MD (figure 8B) can be seen to vary largely along the PD axis. This then presents as a large PD correlation with a small AP correlation (figure 8C - greatly above the unit line). Interestingly, the age-related changes of fneurite_NODDI_ and fneurite_SANDI_ (both approximately representing stick signal fractions) appear to be correlated relatively differently to the positional gradients. fneurite_NODDI_ is more correlated to the PD positional gradient (above the unit line) while fneurite_SANDI_ is more correlated to the AP positional gradient (below the unit line). Analyzing the cosine similarities, it can be seen that the diffusion orientations tend to vary more along AP across age.

**Figure 8.**
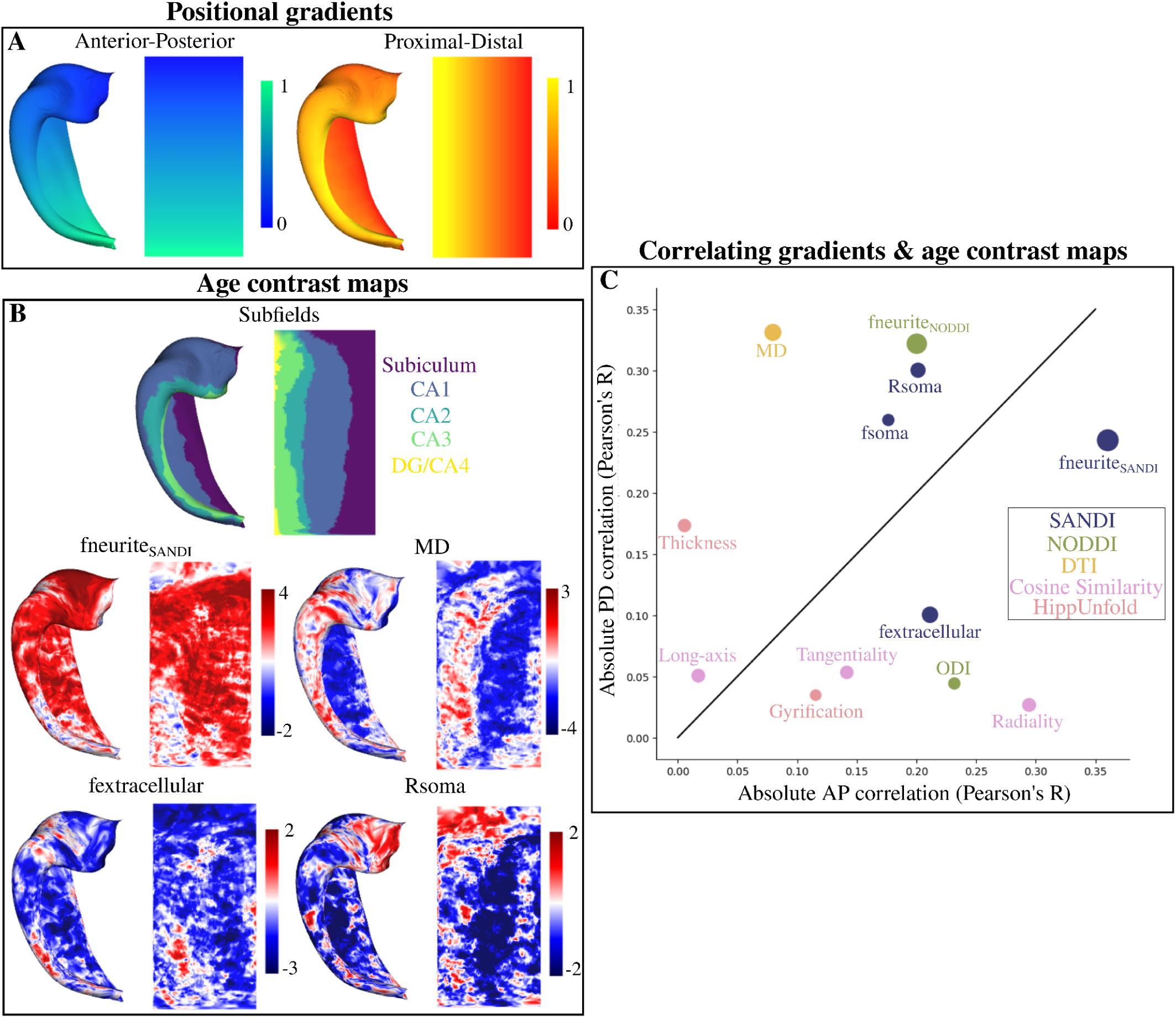
Correlating surface-based age contrasted t-statistic maps with anterior-posterior (AP - long-axis) and proximal-distal (PD - subfields) gradients (DeKraker et al., 2024). (A) Gradients generated on a canonical surface across the AP and PD axis. (B) Subfield parcellation and four examples of a vertex-wise age contrast map with a contrast of age. Thec age contrast maps capture the age-related trends of each metric at each vertex. Maps were calculated with hemisphere-averaged data and plotted on a left hippocampal surface. (C) Absolute correlation (Pearson’s R) of all the age contrast maps (B) with the gradients (A). The size of the points represents the mean of the absolute values of the t-statistic across all vertices - a coarse measure for the total relationship between a metric and age. Colour-coding refers to where each metric is derived from. Identity line is shown as a solid black line.

The correlations between all the age contrast maps (i.e. looking at metric-age covariance) is shown in supplementary figure S6. Overall there appears to be substantial correlation between many of the age-contrasted microstructural maps.

### 2.6 Correlations of vertex-wise microstructural changes across age with MRI, PET, and histology maps

The same age contrast maps in *section 2.5* (figure 9A) were used to correlate with maps derived from histology and MRI at high resolutions (figure 9B) using a hippocampus spin-test (Karat et al., 2023). Figure 9C and the below paragraph presents the correlation and uncorrected p-values between age contrast maps (figure 9A) and all histology/MRI maps (figure 9B).

**Figure 9.**
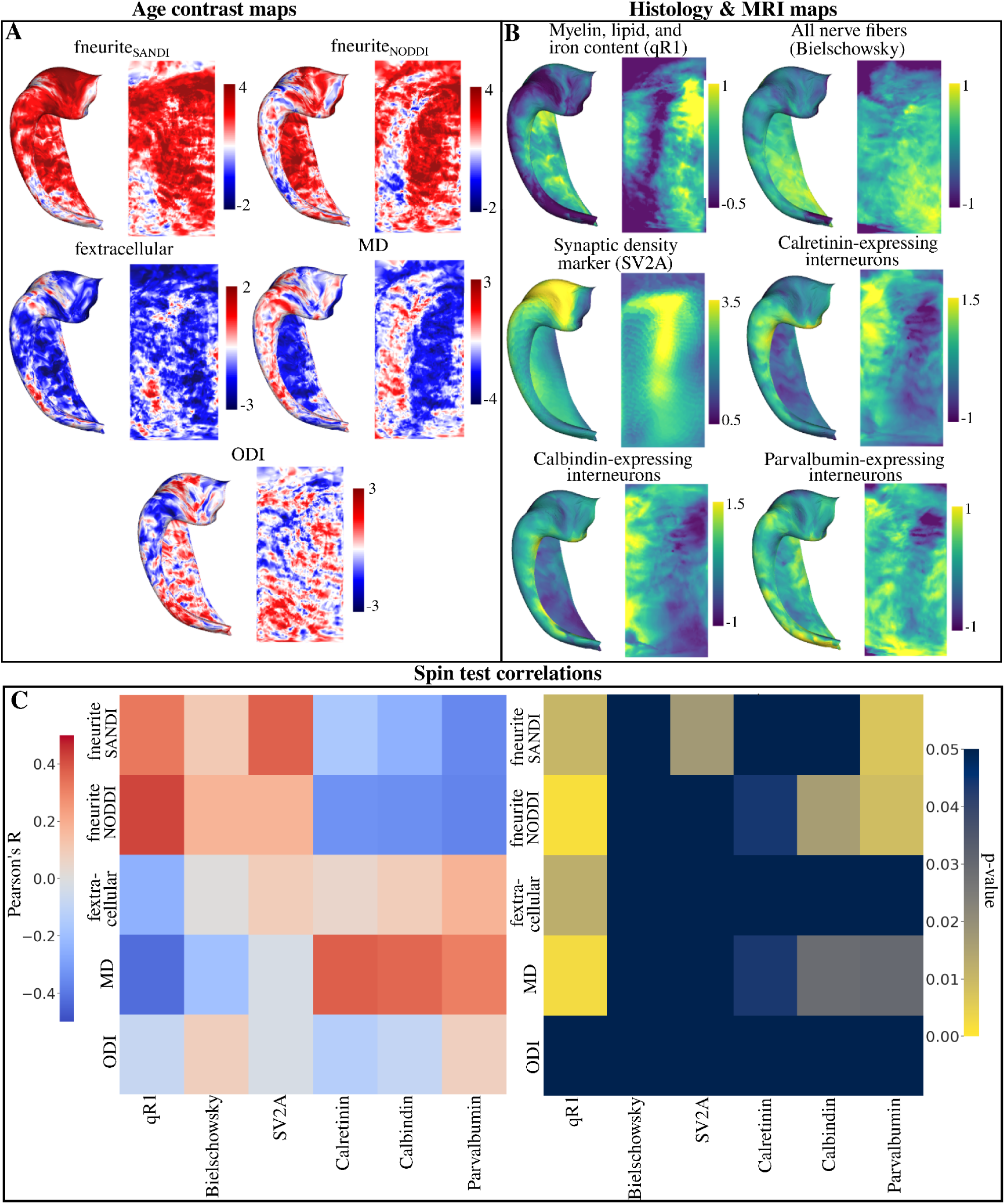
Correlating surface-based age contrast t-statistic maps with maps derived from histology and MRI (DeKraker et al., 2024; Markello et al., 2022). (A) Vertex-wise t-statistic maps with a contrast of age. The age contrast maps capture the age-related trends of each metric at each vertex. Maps were calculated with hemisphere-averaged data and plotted on a left hippocampal surface. (B) Surface maps derived from histological staining and MRI. qR1, Bielschowsky, Calretinin, Calbindin, and Parvalbumin maps were derived from Alkemade et al., 2022. The synaptic vesicle glycoprotein 2A (SV2A) marker was derived from Finnema et al., 2018. (C) Left heatmap displays the Pearson’s R correlation between all age contrast maps in (A) and all histology and MRI maps in (B). Right heatmap displays the uncorrected p-values derived from a hippocampus spin test using 2500 permutations (Karat et al., 2023). Note that the colour bar is inverted such that any brighter component of the heatmap corresponds to a significant p-value.

The age-related changes of fneurite_SANDI_ were positively correlated with qR1 (R=0.33, p=0.01) and SV2A (R=0.37, p=0.017) and negatively correlated with parvalbumin (R=-0.36, p=0.007). The age-related changes of fneurite_NODDI_ were positively correlated with qR1 (R=0.42, p=0.002) and negatively correlated with calretinin (R=-0.34, p=0.04), calbindin (R=-0.34, p=0.02), and parvalbumin (R=-0.36, p=0.007), suggesting further differences between fneurite_SANDI_ and neurite_NODDI_. The fextracellular age contrast map was negatively correlated with qR1 (R=-0.24, p=0.012). Age-related changes in MD were positively correlated with calretinin (R=0.38, p=0.044), calbindin (R=0.36, p=0.029) and parvalbumin (R=0.31, p=0.030) and negatively correlated with qR1 (R=-0.42, p=0.002). Changes to ODI across age were found to not correlate strongly with any histology or MRI map. With FDR correction, only the correlations between fneurite_SANDI_ and qR1, SV2A, and parvalbumin, fneurite_NODDI_ and qR1, calbindin, and parvalbumin, and MD and qR1 remained significant. Finally, we found no correlation between Merker cell body staining and the fsoma age contrast map (R=0.04, p=0.55) and the fsoma map averaged across age (R=0.14, p=0.30) seen in supplementary figure S7.

## 3. Discussion

We probed age-related alterations of hippocampal soma and neurite microstructure using a recent approach for surface-based hippocampal modelling. We found no significant change in volume, gyrification, and thickness across age, while significant changes in microstructure measures related to neurites, soma, and mean diffusivity were found. Sex-specific differences in age-related changes to macro- and microstructure were also found. Leveraging the salient orientation information derived from dMRI, localized diffusion orientation shifts across the hippocampal subfields and long-axis were uncovered which may relate to specific development of the intrahippocampal circuitry. Surface-based analyses suggested large variation in age-related microstructural changes across the proximal-distal (subfield) and anterior-posterior (long-axis) axes, which may reflect ostensible developmental differences along these two axes. Finally, correlation of the age-contrasted microstructure surface maps with MRI, PET, and histology allowed for postulation of the potential underlying microstructure that diffusion changes across age may be sensitive to.

### 3.1 Microstructure changes greatly across age for the subfields and long-axis

Recent evidence has suggested that the hippocampus undergoes a prolonged period of both structural and functional development after birth (Lee et al., 2017). To investigate hippocampal microstructural changes across development, most previous research has used DTI (Basser et al., 1994). In the current study we found MD significantly decreased with age across the subfields and long-axis. At the whole hippocampus level, Callow et al. (2020) found that MD decreased between 4-8 years of age. Langnes et al. (2020) found a protracted period of decreased MD in the anterior hippocampus up until 40 years of age, while posterior MD did not change much in the same range. However, while MD in the current study did appear to change more across age in the anterior portions of the hippocampus, we found the change was also significant in the posterior region. The decrease in MD is likely attributable to an increase in stick-like restrictions, as suggested by the significant increase across age in fneurite_SANDI_ and a corresponding decrease in fextracellular.

We found no significant change in FA from 8-19 years (supplementary figure S8), which conflicts with recent work. Vinci-Booher et al. (2023) found subfield-specific changes in FA across the age range of 5-30 years. In CA1 FA had an apparent parabolic trend across the whole age range. However, the trend of FA in CA1 appeared to decrease linearly when only considering ages 8-19. The DG and subiculum had a reduction in FA across the whole age range, while CA2/3 had an increase in FA (Vinci-Booher et al., 2023). Given that we found no significant change in ODI and a significant increase in fneurite_SANDI_ across age, it may be hypothesized that FA should increase. However, it appears that FA is much more sensitive to changes in ODI than intraneurite signal fractions (figure 10 in Zhang et al., 2012). As well, the range of ODI (0.4-0.6) and fneurite_SANDI_ (0.15-0.30) values found in the current study can be seen to be in a regime of “low” FA contrast (figure 10 in Zhang et al., 2012). Thus it appears that the hippocampal microstructural environment is generally isotropic at 8 years, and the increase in fneurite_SANDI_ is likely coming from an increase in spatially isotropic stick-like restrictions (which ostensibly would not change the already high ODI and thus result in no change in FA). Future investigation of age-related changes in FA is warranted.

Analyzing the primary orientation of diffusion across age, we found an increase in the long-axis oriented diffusion in the anterior lateral and body of the hippocampus with a general corresponding decrease in the radial oriented diffusion. These changes may correspond to the development of intra-hippocampal pathways with known orientation. The fimbria is a white matter pathway which sits atop CA3/CA4 and traverses the hippocampal long-axis where at its most posterior becomes the fornix (Zeineh et al., 2017). The increase in long-axis oriented diffusion which appears to largely occur in the anterior and body of the hippocampus may be capturing changes to the coherent fimbria pathway. However, if this was the case we may also expect to see a decrease in ODI and a corresponding increase in FA in the same region. The increase in long-axis oriented diffusion in these regions could also correspond to a change in the perforant path, which at some levels is oriented across the long-axis (Zeineh et al., 2017).

### 3.2 Microstructure appears much more sensitive to age-related changes then macrostructure

One of the most common metrics used to investigate in vivo hippocampal development is volume (in mm^3^) derived with structural MRI. While common, there are conflicting results related to hippocampal volume changes in childhood and adolescence. We found no change in volume for any long-axis parcel or subfield (measured using HippUnfold and FreeSurfer) between 8-19 years. At the level of the whole hippocampus, some studies have reported an increase in volume in late childhood and adolescence (Krogsrud et al., 2014; Tamnes et al., 2018; Wierenga et al., 2014) while others have found a very slight increase or no volume change in the same age range (Giedd et al., 1996; Uematsu et al., 2012; Coupe et al., 2017; Narvacan et al., 2017; Hu et al., 2013). Coupe et al. (2017) analyzed 2994 subjects across the whole lifespan and found a very fast whole hippocampal volume increase until 8-10 years, followed by a very slow volume increase until 40-50 years. However, the hippocampus is not a monolithic structure, rather, its sub-regions ostensibly have different developmental trajectories. Langnes et al. (2020) found a relative increase in both anterior and posterior hippocampal volumes from 4-20 years while Gogtay et al. (2006) found posterior volumes increased while anterior volumes decreased between 4-25 years. Across all the subfields, Krogsrud et al. (2014) found an increase in volume from ages 4-22 years, with an asymptote occurring around age 16 for all subfields. Contrastingly, Tamnes et al. (2018) found an initial slight increase and then slight decrease in volume in the subiculum and CA1 between ages 8-30. Linear volume decreases were found in CA2/3, CA4, and the DG granular cell layer across the same age range.

Beyond volume, we also analyzed macrostructural measures of both thickness (in mm) and gyrification. Similar to the result for volume, we found no significant change in subfield and long-axis thickness and gyrification acrpss age. This suggests that the general macrostructural form of the hippocampus may stabilize in early childhood. Although, notable differences exist between the current study and previous research (see *section 3.5* for more details). While macrostructure does not appear to change much after 8 years, we showed distinct microstructural changes between 8-19 years suggesting that much of the age-related changes are internal to the hippocampal gray matter (and thus may be invisible to volume measurements). Diffusion in the hippocampus appears to provide improved sensitivity and specificity to hippocampal development then macrostructure (Callow et al., 2020).

### 3.3 Males and females display variable macro- and microstructural trends across age

Previous research has found varying trends in both volume and microstructure between males and females across age. We found that the age-related trends of volume and thickness were significantly different between males and females across the hippocampal subfields. In general, it appeared that females had no significant change in volume and thickness from 8-19 years, while males had an increase in volume and thickness particularly in CA1, CA2, and CA3. This may be explained by previous research which suggests that female hippocampal volumes reach a plateau sooner than male volumes (Pfluger et al., 1999; Uematsu et al., 2012; Hu et al., 2013). However, the evidence for the age at which this plateau occurs (and if it occurs at all) is conflicting. Giedd et al. (1996) found that hippocampal volume increased between ages 4-18 years only in females. Pfluger et al. (1999) found that hippocampal volumes increased much faster in females than in males between 1 month to 15 years of age. That is, at around 2 years of age female hippocampal volumes plateaued, while male volumes kept increasing. This was corroborated by Uematsu et al. (2012) and Hu et al. (2013), where it was shown that females reach their peak hippocampal volume sooner than males. Contrastingly, Tamnes et al. (2018) found no sex differences in hippocampal subregion development between 8-30 years of age, and they found general volume changes in females across the whole age range (i.e. female volumes did not plateau in some subfields).

Few studies have examined sex differences in microstructural development of the hippocampus in late childhood and adolescence. In the current study we found significant sex by age interactions in fneurite_SANDI_ and fextracellular, Rsoma, ODI, and MD, suggesting that the change in microstructure across late childhood and adolescence are different between males and females. Vinci-Booher et al. (2023) found a nonlinear interaction of FA between sex and age in CA2/3. Callow et al. (2020) found sex was not significantly related to hippocampal MD. Interestingly, studies probing glia and neuron density and morphology in the Macaque hippocampus found no significant differences between males and females from juvenile to geriatric age, or from birth up to 1-year (Robillard et al., 2016; Jabès et al., 2010). However, these studies were limited to small sample sizes which makes any strong conclusion difficult. Future research is needed to further understand the developmental differences of hippocampal microstructure between males and females in the formative period of late childhood and adolescence.

### 3.4 Age-related microstructural changes correlate to hippocampal axes and specific histological metrics

The hippocampus is generally studied in the context of subfields which are the structurally distinct subunits of the hippocampus (Karat et al., 2024; Ding and Van Hoesen, 2015). Recently the hippocampal long-axis (anterior to posterior) has garnered substantial interest given evidence that it is also structurally and functionally distinct (Chase et al., 2015; Strange et al., 2014; Nichols et al., 2023; Poppenk et al., 2013). Here we investigated the age-related changes in microstructure within the context of the subfields and long-axis at the vertex level. We showed that age-related changes in MD were much more correlated with the proximal-distal (i.e., subfields) than the long-axis, suggesting that the subfields have greater age-related variability from the perspective of diffusivity. Previous research has found strong associations of age with MD particularly in the subiculum and CA1, where it does appear that age-related changes in MD vary more across the subfields than the long-axis (Wolf et al., 2015). However, it has also been shown that the anterior hippocampus has an extended period of MD changes across late childhood and adolescence, while the posterior hippocampus remained relatively unchanged (Langnes et al., 2020). Interestingly, age-related changes in fneurite_SANDI_ varied more across the long-axis than the subfields, while Rsoma and fsoma showed trends similar to MD.

To further contextualize the results, we correlated MRI, PET, and histology maps of specific microstructural features with the age-related diffusion microstructure maps. Interestingly, we found that age-related changes in fneurite_SANDI_ correlated with qR1 (myelin, lipids, iron content) and a marker of synaptic density, SV2A. Put differently, in regions of high qR1/synaptic density, fneurite_SANDI_ increased across age. Similarly, we found that the age-related changes in MD correlated with qR1, such that in regions of high qR1 MD decreased with age, and vice versa. The age-related changes in fneurite_SANDI_, fextracellular and MD thus may be related to myelin alterations in the hippocampus. Previous research has shown that even at 11 years of age the density of myelinated fibers in humans did not reach adult levels, suggesting myelination continues through and beyond late childhood/adolescence (Ábrahám et al., 2010). Similarly, recent work has shown that oligodendrocyte-specific gene expression increased with age, indicating subsequent myelination processes (Genc et al., 2024). Occurring concurrently with myelination is apparent glia alterations. In Macaques it was found that astrocyte process length and complexity increased from juvenile to adulthood (Robillard et al., 2016). An increase in stick-like glia processes across age may also ostensibly change fneurite_SANDI_. As well, the correlation of SV2A with fneurite_SANDI_ may be a result of increased dendrite ramifications. Mellström et al. (2016) found a quadratic relationship between spine density and branching order of the basal dendrites of CA1 pyramidal neurons, suggesting that as branching order increases (more “sticks” from the diffusion perspective) there is higher spine density, and thus a greater synaptic density. Finally, in the current study we found that fsoma did not change much across age, and that it was not correlated to a Merker stain for cell bodies. Indeed, Jabès et al., (2010) found that the number of principal neurons in the Macaque hippocampus did not change significantly from birth to 5-9 years of age in all subfields except the granule cell layer of the DG, which we did not have the resolution to capture in the current study. However, it should be noted that fsoma may not be expected to correlate with a stain for cell bodies given its a signal fraction rather than a volume fraction, as it does not account for T2 differences between microstructural compartments. The apparent decrease in Rsoma across age could be potentially related to an increase in glia presence. Indeed, gene expression analyses in Genc et al. (2024) suggests that across age there is a decrease in cells with larger soma (i.e., endothelial) which are outweighed by an increase in cells with smaller soma (i.e., oligodendrocytes).

### 3.5 Limitations

One limitation of the current work was the 2mm isotropic diffusion image resolution. Given the thickness of the hippocampus, some partial voluming with surrounding CSF and extra-hippocampal WM was expected. However, we attempted to minimize partial voluming by sampling the diffusion measures along the middle of the hippocampal GM. To examine how effective this was, we used FSL’s FAST tool to generate tissue type probabilities at the higher T1w resolution. We found relatively low CSF and WM (apart from the subiculum) tissue probability, suggesting that the sampling across the midthickness surface was mostly within the GM. Another limitation was the correlation of the age-related microstructure t-statistic maps with static maps derived from adult histology. Ideally the comparison would be done with histology-derived age contrasted t-statistic maps capturing the cross-sectional changes in a particular histological measure across development. As well, the current study had a moderate sample size at 88 participants with a mean age of 12.6 years. That is, there were more samples in the younger 8-12 year age range then the older 14-19 year range which could make the results more reflective of changes occurring mainly in the late childhood stage. Given some of the strong correlations found in the current study, it would be valuable to examine the same advanced diffusion metrics across the whole lifespan instead of the narrower slice of childhood and adolescence. Likewise, it would be useful to probe diffusion changes longitudinally between childhood and adolescence, rather than the cross-sectional design used here. Finally, there is inherent variability in contemporary methods which seek to provide measures of hippocampal subfield volume through segmentation (HippUnfold, FreeSurfer, ANTs, manual delineation, etc…). Given this variability it can be difficult to compare results pertaining to subfield and long-axis volume across studies. However, in the current study both HippUnfold and FreeSurfer provided converging evidence that hippocampal volume did not change.

## 4. Conclusion

The hippocampus serves multiple cognitive functions, yet little is known about its microstructural development. Here we report, for the first time, distinct neurite and soma developmental profiles in the hippocampus during late childhood and adolescence using advanced diffusion modelling. Specifically, we report an age-related increase in neurite fraction and concurrent decrease in extracellular fraction and soma radius which appears to be subfield and long-axis specific. Future research should look to examine the same diffusion measures across the whole lifespan, and correlate these with cognitive processes that rely on the hippocampus such as memory and spatial navigation.

## 5. Materials and methods

### 5.1 Data acquisition and preprocessing

88 participants aged 8-19 years (42 male, mean age=12.6, SD=2.9) were scanned on a 3T Siemens Connectom system with ultra-strong (300 mT/m) gradients (Genc et al., 2020; Genc et al., 2024). Structural T1-weighted (T1w) images were acquired at 1mm isotropic resolution (TE=2 ms, TR=2300 ms). Diffusion MRI data were acquired at 2 mm isotropic resolution (TE=59 ms, TR=3000 ms) with b-values of 0 (14 volumes, interleaved), 0.5 (30 directions), 1.2 (30 directions), 2.4 (60 directions), 4.0 (60 directions), and 6.0 (60 directions) ms/µm^2^. Diffusion directions were determined using an electrostatic repulsion algorithm generalized across all shells (Caruyer et al., 2013). Data were acquired using an anterior-posterior phase-encoding direction, with one additional inverse phased encoding (posterior-anterior) volume. The total dMRI acquisition time was 16 minutes and 14 seconds.

Data pre-processing has been described previously in Genc et al. (2020). Briefly, preprocessing steps included: denoising (Veraart et al., 2016), slice-wise outlier detection (Sairanen et al., 2018), drift correction (Vos et al., 2016), motion, eddy, and susceptibility-induced distortion correction (Andersson & Sotiropoulos, 2016), Gibbs ringing correction (Kellner et al., 2016), bias field correction (Tustison et al., 2010), and gradient non-uniformity correction (Rudrapatna et al., 2018).

### 5.2 Surface modelling with HippUnfold

An automated software (HippUnfold) for hippocampal subfield segmentation and surface-based mapping was used in this study (DeKraker et al., 2022). HippUnfold uses a deep neural network (a ‘U-net’; Isensee et al., 2021) to segment within each subject the hippocampal gray matter, dentate gyrus (DG), stratum radiatum lacunosum moleculare (SRLM), and the topological bounds of the hippocampus, including the hippocampal-amygdala transition area, medial temporal lobe cortex, pial surface, and the indusium griseum. Using the gray matter as the domain of interest and the previously defined hippocampal boundaries, Laplace coordinates are generated along the anterior-posterior, proximal-distal, and inner-outer directions which define a complete 3D coordinate system of the hippocampus which respects subject morphology. Using these coordinates, it is then possible to project subfields defined from a histological atlas to each subject in their native space, as well as generate surfaces at varying inner-outer/laminar depths. The subfields include the subiculum, Cornu Ammonis (CA) 1-4, the DG, and the SRLM. In the current work, the DG and CA4 were averaged together. As well, using the above coordinates and generated surfaces we parcellated the hippocampal anterior-posterior (also referred to as the long-axis) into 5 bins including the anterior uncus, anterior lateral, body anterior, body posterior, and posterior/tail. For more detailed methods see DeKraker et al. (2022).

All segmentations were reviewed by author BGK for errors. Of the 88 subjects, 14 U-net segmentations were manually corrected for small over- or underestimations of tissue, and HippUnfold was rerun using the manually corrected segmentations. The metrics from HippUnfold used in this study were gyrification and thickness which are calculated on the generated surfaces, and the more traditional measure of subfield volume calculated in each subject’s native space. All other volumetric measures of interest (i.e., the microstructure maps) were sampled onto the midthickness (middle of the gray matter) surface for visualization and analysis. All midthickness surfaces used in this study were composed of 7262 vertices, roughly corresponding to a spacing of 0.5 mm. Connectome Workbench (https://github.com/Washington-University/workbench) was used to sample values at each surface vertex from volume data using the enclosing method. To investigate potential partial volume effects PVE, FSL’s FAST tool was used to generate partial volume tissue estimates of gray matter (GM), white matter (WM), and cerebrospinal fluid (CSF) using the T1w images (Zhang et al., 2001). These partial volume maps were then linearly interpolated to the lower 2 mm isotropic diffusion resolution and sampled on the midthickness surface in the same way as the other metrics of interest. Analyses were then performed to examine if estimated tissue type probabilities of GM, WM, and CSF varied with age. Finally, to provide a complementary measure of subfield volume, FreeSurfer (version 7.2.0) was run across all subjects.

### 5.3 Microstructural modelling

The diffusion MRI data were analyzed using DTI (Basser et al., 1994), NODDI (Zhang et al., 2012), and SANDI (Palombo et al., 2020).

The FMRIB Software Library (FSL; version 6.0.5, Smith et al., 2004) was used to fit the diffusion tensor using all b = 0, b = 0.5, and b = 1.2 ms/µm^2^ data. DTI characterizes the diffusion process as a symmetric 3×3 tensor which describes diffusion in different directions, where the diffusion signal can be written as:

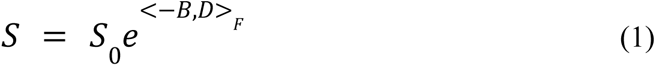

where S is the diffusion signal along a particular direction, B is the b-matrix (Mattiello et al., 1997), D is the diffusion tensor, and <*>_F_ is the Frobenius inner product. The eigendecomposition of D provides useful metrics which capture ensemble diffusion characteristics, including fractional anisotropy (FA; the variance of the eigenvalues - preferential diffusion along a particular direction) and the mean diffusivity (MD; the mean of the eigenvalues - average diffusivity in physical units). While DTI has been extensively used to interrogate brain microstructure, the complex nature of water diffusion in neural tissue can not usually be well described by a single tensor (Jones et al., 2013).

The Microstructure Diffusion Toolbox (MDT; version 1.2.7) (Harms et al., 2017) was used to fit the NODDI model using all b-values (Zhang et al., 2012). The NODDI model assumes that water diffusion is occurring in three non-exchanging microstructural environments consisting of intraneurite, extraneurite, and CSF compartments. The intraneurite compartment generally refers to the space enclosed by the membrane of neurites (assumed impermeable to water) and is modelled as a set of orientation dispersed sticks (i.e., cylinders with zero radius) according to the Watson distribution (Zhang et al., 2012). The extraneurite compartment comprises the space around neurites and is modelled as a cylindrically symmetric tensor. The parallel diffusivity of the intraneurite and extraneurite compartment was fixed to 1.7 µm^2^/ms. The CSF compartment is modelled as isotropic gaussian diffusion described by a single diffusion coefficient which was fixed to 3.0 µm^2^/ms. The diffusion signal according to the NODDI model can be written as (Zhang et al., 2012):

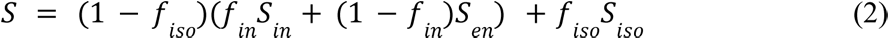

where *f_iso_, f_in_, f_en_*, is the signal fraction from the CSF, intraneurite, and extraneurite compartments respectively, while *S_iso_, S_in_, S_en_* is the diffusion signal arising from the CSF, intraneurite, and extraneurite compartments respectively. The two metrics from NODDI investigated in this study include the *f_in_* which will be referred to as fneurite_NODDI_ and the orientation dispersion index (ODI) which is derived from the Watson distribution.

To fit the SANDI model, the SANDI matlab toolbox (https://github.com/palombom/SANDI-Matlab-Toolbox-v1.0) (Palombo et al., 2020; Ianuş et al., 2022) was used. The SANDI model assumes that water diffusion occurs in the non-exchanging intraneurite, extracellular, and intrasoma compartments. Much like the NODDI model, the intraneurite compartment is represented as diffusion within sticks. The extracellular compartment is modelled as isotropic gaussian diffusion characterized by a single diffusion coefficient. Finally, the intrasoma compartment is modelled as diffusion occurring in a restricting sphere with a radius *r*_s_. The diffusion signal according to the SANDI model can be written as (Palombo et al., 2020):

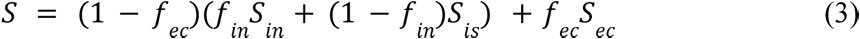

where *f*_*ec*_ and *f*_*in*_ are the signal fractions from the extracellular and intraneurite compartments, respectively, and *f*_*is*_ = (1 − *f*_*in*_) is the signal fraction of the soma compartment. *S*_*ec*_, *S*_*in*_, *S*_*is*_ are the diffusion signals arising from the extracellular, intraneurite, and soma compartments, respectively. Beyond the addition of the soma compartment, the SANDI model focuses on estimating microstructural features that are orientation independent. That is, the diffusion signal is averaged across all gradient directions before model fitting. Thus the direction-averaged formulations of *S*_*ec*_ and *S*_*in*_ depart from the functional forms of the signals in NODDI which have an orientation dependence. Furthermore, unlike NODDI, SANDI fits the diffusivities of the intraneurite and extracellular compartments, while fixing the intrasoma diffusivity to 3.0 µm^2^/ms. The maps from SANDI analyzed in this study include fextracellular = *f*_*ec*_ , fneurite_SANDI_ = (1 − *f*_*ec*_)*f*_*in*_; fsoma = (1 − *f*_*ec*_)(1 − *f*_*in*_) and *r*_*s*_ referred to as Rsoma.

Diffusion data and the corresponding metric maps were registered to each subject’s T1w space. Once aligned, all metrics of interest were sampled on the hippocampal midthickness surface as described in *section 5.2*. From the surface maps, metrics were then averaged within subfields or along the anterior-posterior axis for further analysis.

### 5.4 Orientation cosine similarities

Microstructure in the hippocampus tends to be aligned stereotypically along the anterior-posterior (AP; long-axis), proximal-distal (PD; tangential), and/or inner-outer (IO; radial) axes (Duvernoy et al., 2013; Nieuwenhuys et al., 2008; Zeineh et al., 2017; Karat et al., 2023). Diffusion analysis can provide a fiber orientation distribution function (fODF), where its peaks ostensibly correspond to the orientation of the underlying microstructure (Jeurissen et al., 2014). Similar to Karat et al. (2023), analyses were performed to examine how the first peak of the fODF was oriented relative to the three hippocampal axes, and how these orientations may shift across age. To calculate the fODF, we used the MRtrix3 toolbox (version 3.0.3) (Tournier et al., 2019) and multi-shell multi-tissue constrained spherical deconvolution (MSMT-CSD) (Jeurissen et al., 2014) using all b-values with a response function averaged across all subjects. We then extracted the first/largest peak from the spherical harmonic representation of the WM fODF (i.e., the orientation with the maximum signal change). To get an orientation measure along the three hippocampal axes, we obtained gradient vector fields along the AP, PD, and IO hippocampal axes by taking the first partial derivative of the respective Laplacian coordinates provided by HippUnfold along the x, y, and z spatial dimensions (Karat et al., 2023):

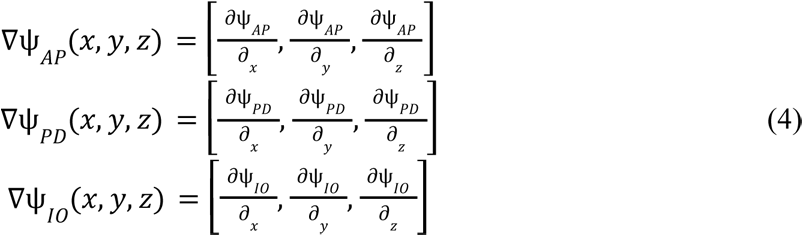

where ψ_*_ is the Laplacian scalar field along one hippocampal axis which was obtained by solving Laplace’s equation: 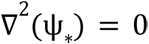 (DeKraker et al., 2022). The result of equation 4 is three vector images for each hemisphere and subject, which point along a single hippocampal axis (AP, PD, or IO). With the definition of the inner product between two vectors as *u* . *v* = |*u*||*v*|*cos*(θ), we take these vector fields and the aligned first peak of the fODF and calculate voxel-wise cosine similarities as:

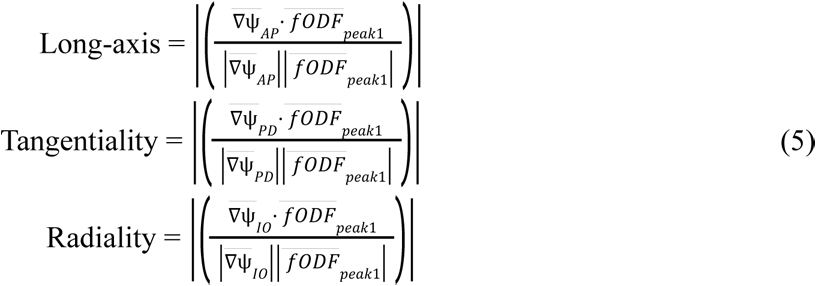

which is depicted in the top of figure 6. A cosine similarity of 0 corresponds to a case where the two vectors are orthogonal, while a cosine similarity of 1 corresponds to a case where the vectors are parallel. A higher cosine similarity thus relates to a case where diffusion is highly oriented along a particular hippocampal axis.A total of three scalar cosine similarity metrics were generated for each subject and hemisphere, and these metrics were sampled along the midthickness surface as described above.

### 5.5 Statistics, correlations, and histology mapping

Multiple statistics were used to examine macro- and microstructure across age, sex, and hemisphere.

#### 5.5.1 Correlations between age and averaged macro- and microstructure

Pearson’s correlation coefficient was used to examine the relation between age and all macro- and microstructural measures at the subfield and anterior-posterior averaged level. Using the pearsonr function in Scipy (version 1.11.3), hypothesis testing was performed to determine the probability that two uncorrelated variables could produce a correlation similar to the observed correlation value. Given the large number of exploratory correlations performed, we sought to set a minimum alpha for which the Pearson’s R would be considered significant. We set the minimum alpha to 0.01, which is analogous to a Bonferroni correction corresponding to 5 hypothesis tests (testing a metric across the 5 subfields). We also report thresholds of 0.005 and 0.0005 so that the correlations can be considered under a more conservative correction.

#### 5.5.2 Interactions between sex, hemisphere, and age

General linear modelling was used to examine the significance of the interaction between sex and age and hemisphere and age. That is, we sought to determine if the slopes of the macro- and microstructure metrics across age were significantly different between sexes and hemispheres. First a full linear model was built containing all relevant terms, then a reduced model was built which contained all the same terms but removed a single variable of interest (sex and age or hemisphere and age interaction). An F-test was then performed between the full and reduced model to examine if the variable of interest resulted in a significantly better model (i.e. it captured a significant portion of variance in the dependent variable (DV)):

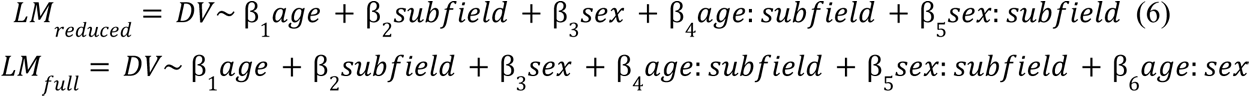

where : implies an interaction between two variables and the DV was a given macro- or microstructural metric. To examine anterior-posterior differences, similar models were built by replacing the subfield term with an anterior-posterior parcellation term instead. Note that for brevity not all β coefficients were shown here since each subfield and its interactions required its own coefficients. In this example both models were identical except for an additional interaction term between age and sex. An F-test between the reduced and full model was then used to examine if the age by sex interaction term significantly improved the linear model. A significant F-test suggested that the change in the DV across age was significantly different between males and females. False-discovery rate (FDR) correction was then applied to the p-values derived from the F-test. The same analysis was performed using an interaction of age and hemisphere, as well as age and subfield.

#### 5.5.3 Vertex-wise statistics

The statistics mentioned thus far were applied only to subfield and anterior-posterior averaged values. To provide increased spatial fidelity of any age-related macro- or microstructural change, statistics were also performed at the vertex level. At each vertex on the hippocampal surface a linear model was built of the form:

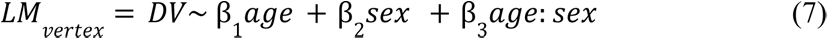

where the t-statistic for the contrast of age was extracted (where *t*_*age*_ = β_1_ /*SD* where *SD* is the standard deviation of β_1_ ). These age contrasted t-statistic maps (referred to as age contrast maps) capture the linear age-related changes of each metric at each vertex. The age contrast maps were then correlated with contrived anterior-posterior and proximal-distal positional gradient maps using Pearson’s R (Figure 8A). The correlations with the gradients describe along which hippocampal axis are potential age-related microstructural changes occurring.

#### 5.5.4 Correlations with MRI, PET, and histology

The age contrast maps (described in *section 5.5.3*) were correlated with high resolution MRI, PET, and histology maps using a hippocampus spin test (Karat et al., 2023). All metrics described below were sampled along the midthickness surface of the hippocampus. Maps of quantitative R1 (qR1) were averaged across 4 ex vivo hippocampi at 9.4T MRI and are thought to capture myelin, lipid, and iron content (DeKraker et al., 2024; Lutti et al., 2014; Stüber et al., 2014). Bielschowsky histological staining was averaged across 4 hippocampi and is representative of all types of nerve fibers (Uchihara, 2007; Alkemade et al., 2022; DeKraker et al., 2024). Synaptic vesicle glycoprotein 2A (SV2A) PET maps which are representative of synaptic density were averaged across 76 in vivo hippocampi (Rossi et al., 2022). Merker histological staining was averaged across 4 hippocampi and captures neuronal cell bodies (DeKraker et al., 2024; Amunts et al., 2013). Finally, interneuron histological markers of calbindin, calretinin, and parvalbumin were averaged across 4 hippocampi (Alkemade et al., 2022; DeKraker et al., 2024). Given the large number of correlations here, we focus on the uncorrected p-values for discussion (although we still report the FDR-corrected p-values).

## Acknowledgements and funding sources

The data were acquired at the UK National Facility for In Vivo MR Imaging of Human Tissue Microstructure funded by the EPSRC (grant EP/M029778/1) and The Wolfson Foundation. BGK is supported by a post-graduate scholarship from the Natural Sciences and Engineering Research Council of Canada (NSERC). SG is supported by the Royal Children’s Hospital Foundation (RCHF 2022-1402). EPR is supported by NICHD at NIH (F32HD103313). MP is funded by UKRI Future Leaders Fellowship (MR/T020296/2). ARK is supported by the Canada Research Chairs program #950-231964, NSERC Discovery Grants RGPIN-2015-06639 and RGPIN-2023-05558, Canadian Institutes for Health Research Project grant #366062, Canada Foundation for Innovation (CFI) John R. Evans Leaders Fund project #37427, the Canada First Research Excellence Fund, and Brain Canada. DKJ is supported by a Wellcome Trust Investigator Award (096646/Z/11/Z) and a Wellcome Trust Strategic Award (104943/Z/14/Z).

We thank the children/adolescents (and their carers) for generously donating their time to participate in this study.

## References

Ábrahám, H., Vincze, A., Jewgenow, I., Veszprémi, B., Kravják, A., Gömöri, É., & Seress, L. (2010). Myelination in the human hippocampal formation from Midgestation to adulthood. International Journal of Developmental Neuroscience, 28(5), 401–410.

Amunts, K., Lepage, C., Borgeat, L., Mohlberg, H., Dickscheid, T., Rousseau, M.-É., Bludau, S., Bazin, P.-L., Lewis, L. B., Oros-Peusquens, A.-M., Shah, N. J., Lippert, T., Zilles, K., & Evans, A. C. (2013). BigBrain: An ultrahigh-resolution 3D human brain model. Science, 340(6139), 1472–1475.

Alkemade, A., Bazin, P.-L., Balesar, R., Pine, K., Kirilina, E., Möller, H. E., Trampel, R., Kros, J. M., Keuken, M. C., Bleys, R. L., Swaab, D. F., Herrler, A., Weiskopf, N., & Forstmann, B. U. (2022). A unified 3D map of microscopic architecture and MRI of the human brain. Science Advances, 8(17).

Andersson, J. L. R., & Sotiropoulos, S. N. (2016). An integrated approach to correction for off-resonance effects and subject movement in diffusion MR imaging. NeuroImage, 125, 1063–1078.

Basser, P. J., Mattiello, J., & LeBihan, D. (1994). Mr diffusion tensor spectroscopy and imaging. Biophysical Journal, 66(1), 259–267.

Buzsáki, G., & Moser, E. I. (2013). Memory, navigation and Theta rhythm in the hippocampal-entorhinal system. Nature Neuroscience, 16(2), 130–138.

Callow, D. D., Canada, K. L., & Riggins, T. (2020). Microstructural integrity of the hippocampus during childhood: Relations with age and source memory. Frontiers in Psychology, 11.

Caruyer, E., Lenglet, C., Sapiro, G., & Deriche, R. (2013). Design of multishell sampling schemes with uniform coverage in diffusion MRI. Magnetic Resonance in Medicine, 69(6), 1534–1540.

Chase, H. W., Clos, M., Dibble, S., Fox, P., Grace, A. A., Phillips, M. L., & Eickhoff, S. B. (2015). Evidence for an anterior–posterior differentiation in the human hippocampal formation revealed by meta-analytic parcellation of fMRI coordinate maps: Focus on the subiculum. NeuroImage, 113, 44–60.

Coupé, P., Catheline, G., Lanuza, E., & Manjón, J. V. (2017). Towards a unified analysis of brain maturation and aging across the entire lifespan: A MRI analysis. Human Brain Mapping, 38(11), 5501–5518.

DeKraker, J., Haast, R. A., Yousif, M. D., Karat, B., Lau, J. C., Köhler, S., & Khan, A. R. (2022). Automated hippocampal unfolding for morphometry and subfield segmentation with Hippunfold. eLife, 11.

DeKraker, J., Cabalo, D., Royer, J., Khan, A., Karat, B., Benkarim, O., Cruces, R. R., Frauscher, B., Pana, R., Hansen, J., Misic, B., Valk, S., Lau, J., Kirschner, M., Bernsconi, A., Bernasconi, N., Muenzing, S., Axer, M., Amunts, K., … Bernhardt, B. (2024). HippoMaps: Multiscale Cartography of Human Hippocampal Organization. bioRxiv.

Ding, S., & Van Hoesen, G. W. (2015). Organization and detailed parcellation of human hippocampal head and body regions based on a combined analysis of cyto- and chemoarchitecture. Journal of Comparative Neurology, 523(15), 2233–2253.

Duvernoy, H. M., Cattin, F., & Risold, P. Y. (2013). The human hippocampus: Functional anatomy, vascularization, and serial sections with MRI. Springer.

Finnema, S. J., Nabulsi, N. B., Mercier, J., Lin, S., Chen, M.-K., Matuskey, D., Gallezot, J.-D., Henry, S., Hannestad, J., Huang, Y., & Carson, R. E. (2018). Kinetic Evaluation and test–retest reproducibility of [11C]UCB-J, a novel radioligand for positron emission tomography imaging of synaptic vesicle glycoprotein 2a in humans. Journal of Cerebral Blood Flow & Metabolism, 38(11), 2041–2052.

Genc, S., Tax, C. M., Raven, E. P., Chamberland, M., Parker, G. D., & Jones, D. K. (2020). Impact of *b*-value on estimates of apparent fibre density. Human Brain Mapping, 41(10), 2583–2595.

Genc, S., Ball, G., Chamberland, M., Raven, E. P., Tax, C. M., Ward, I., Yang, J. Y.-M., Palombo, M., & Jones, D. J. K. (2024). MRI Signatures of Cortical Microstructure in Human Development Align with Oligodendrocyte Cell-Type Expression. bioRxiv.

Giaccio, R. G. (2006). The dual origin hypothesis: An evolutionary brain-behavior framework for analyzing psychiatric disorders. Neuroscience & Biobehavioral Reviews, 30(4), 526–550.

Giedd, J. N., Vaituzis, A. C., Hamburger, S. D., Lange, N., Rajapakse, J. C., Kaysen, D., Vauss, Y. C., & Rapoport, J. L. (1996). Quantitative MRI of the temporal lobe, amygdala, and hippocampus in normal human development: Ages 4-18 years. The Journal of Comparative Neurology, 366(2), 223–230.

Gogtay, N., Nugent, T. F., Herman, D. H., Ordonez, A., Greenstein, D., Hayashi, K. M., Clasen, L., Toga, A. W., Giedd, J. N., Rapoport, J. L., & Thompson, P. M. (2006). Dynamic mapping of normal human hippocampal development. Hippocampus, 16(8), 664–672.

Goldman-Rakic, P. S. (1987). Development of cortical circuitry and cognitive function. Child Development, 58(3), 601.

Harms, R. L., Fritz, F. J., Tobisch, A., Goebel, R., & Roebroeck, A. (2017). Robust and fast nonlinear optimization of diffusion mri microstructure models. NeuroImage, 155, 82–96.

Hu, S., Pruessner, J. C., Coupé, P., & Collins, D. L. (2013b). Volumetric analysis of medial temporal lobe structures in brain development from childhood to adolescence. NeuroImage, 74, 276–287.

Ianuş, A., Carvalho, J., Fernandes, F. F., Cruz, R., Chavarrias, C., Palombo, M., & Shemesh, N. (2022). Soma and neurite density MRI (Sandi) of the in-vivo mouse brain and comparison with the allen brain atlas. NeuroImage, 254, 119135.

Isensee, F., Jaeger, P. F., Kohl, S. A., Petersen, J., & Maier-Hein, K. H. (2021). NNU-net: A self-configuring method for deep learning-based biomedical image segmentation. Nature Methods, 18(2), 203–211.

Jabès, A., Lavenex, P. B., Amaral, D. G., & Lavenex, P. (2010). Postnatal development of the Hippocampal Formation: A stereological study in Macaque Monkeys. The Journal of Comparative Neurology, 519(6), 1051–1070.

Jelescu, I. O., & Budde, M. D. (2017). Design and validation of diffusion MRI models of white matter. Frontiers in Physics, 5.

Jensen, J. H., & Helpern, J. A. (2010). MRI quantification of non-gaussian water diffusion by kurtosis analysis. NMR in Biomedicine, 23(7), 698–710.

Jeurissen, B., Tournier, J.-D., Dhollander, T., Connelly, A., & Sijbers, J. (2014). Multi-tissue constrained spherical deconvolution for improved analysis of multi-shell diffusion MRI data. NeuroImage, 103, 411–426.

Jones, D. K., Alexander, D. C., Bowtell, R., Cercignani, M., Dell’Acqua, F., McHugh, D. J., Miller, K. L., Palombo, M., Parker, G. J. M., Rudrapatna, U. S., & Tax, C. M. W. (2018). Microstructural imaging of the human brain with a ‘super-scanner’: 10 key advantages of ultra-strong gradients for diffusion MRI. NeuroImage, 182, 8–38.

Jones, D. K., Knösche, T. R., & Turner, R. (2013). White matter integrity, fiber count, and other fallacies: The do’s and don’ts of Diffusion MRI. NeuroImage, 73, 239–254.

Karat, B. G., DeKraker, J., Hussain, U., Köhler, S., & Khan, A. R. (2023). Mapping the macrostructure and microstructure of the in vivo human hippocampus using diffusion MRI. Human Brain Mapping, 44(16), 5485–5503.

Karat, B. G., Kohler, S., & Khan, A. R. (2024). Diffusion MRI of the hippocampus. Journal of Neuroscience, 44(23).

Kellner, E., Dhital, B., Kiselev, V. G., & Reisert, M. (2016). Gibbs-ringing artifact removal based on local subvoxel-shifts. Magnetic Resonance in Medicine, 76, 1574–1581.

Krogsrud, S. K., Tamnes, C. K., Fjell, A. M., Amlien, I., Grydeland, H., Sulutvedt, U., Due-Tønnessen, P., Bjørnerud, A., Sølsnes, A. E., Håberg, A. K., Skrane, J., & Walhovd, K. B. (2014). Development of hippocampal subfield volumes from 4 to 22 years. Human Brain Mapping, 35(11), 5646–5657.

Langnes, E., Sneve, M. H., Sederevicius, D., Amlien, I. K., Walhovd, K. B., & Fjell, A. M. (2020). Anterior and posterior hippocampus macro- and microstructure across the lifespan in relation to memory—a longitudinal study. Hippocampus, 30(7), 678–692.

Lebel, C., Walker, L., Leemans, A., Phillips, L., & Beaulieu, C. (2008). Microstructural maturation of the human brain from childhood to adulthood. NeuroImage, 40(3), 1044–1055.

Lee, J. K., Ekstrom, A. D., & Ghetti, S. (2014). Volume of hippocampal subfields and episodic memory in childhood and adolescence. NeuroImage, 94, 162–171.

Lee, J.K., Johnson, E.G., Ghetti, S. (2017). Hippocampal Development: Structure, Function and Implications. In: Hannula, D., Duff, M. (eds) The Hippocampus from Cells to Systems. Springer, Cham.

Le Bihan, D. (1995). Molecular diffusion, tissue microdynamics and microstructure. NMR in Biomedicine, 8(7), 375–386.

Lutti, A., Dick, F., Sereno, M. I., & Weiskopf, N. (2014). Using high-resolution quantitative mapping of R1 as an index of cortical myelination. NeuroImage, 93, 176–188.

Markello, R. D., Hansen, J. Y., Liu, Z.-Q., Bazinet, V., Shafiei, G., Suárez, L. E., Blostein, N., Seidlitz, J., Baillet, S., Satterthwaite, T. D., Chakravarty, M. M., Raznahan, A., & Misic, B. (2022). Neuromaps: Structural and Functional Interpretation of Brain Maps.

Mattiello, J., Basser, P. J., & Le Bihan, D. (1997). The B matrix in diffusion tensor echo-planar imaging. Magnetic Resonance in Medicine, 37(2), 292–300.

McNab, J. A., Edlow, B. L., Witzel, T., Huang, S. Y., Bhat, H., Heberlein, K., Feiweier, T., Liu, K., Keil, B., Cohen-Adad, J., Tisdall, M. D., Folkerth, R. D., Kinney, H. C., & Wald, L. L. (2013). The Human Connectome Project and Beyond: Initial applications of 300MT/m gradients. NeuroImage, 80, 234–245.

Mellström, B., Kastanauskaite, A., Knafo, S., Gonzalez, P., Dopazo, X. M., Ruiz-Nuño, A., Jefferys, J. G., Zhuo, M., Bliss, T. V., Naranjo, J. R., & DeFelipe, J. (2016). Specific cytoarchitectureal changes in hippocampal subareas in Dadream Mice. Molecular Brain, 9(1).

Narvacan, K., Treit, S., Camicioli, R., Martin, W., & Beaulieu, C. (2017). Evolution of deep gray matter volume across the human lifespan. Human Brain Mapping, 38(8), 3771–3790.

Nichols, E. S., Blumenthal, A., Kuenzel, E., Skinner, J. K., & Duerden, E. G. (2023). Hippocampus long-axis specialization throughout development: A meta-analysis. Human Brain Mapping, 44(11), 4211–4224.

Nieuwenhuys, R., van Huijzen, C., & Voogd, J. (2008). The human central nervous system. Springer.

Palombo, M., Ianus, A., Guerreri, M., Nunes, D., Alexander, D. C., Shemesh, N., & Zhang, H. (2020). SANDI: A compartment-based model for non-invasive apparent Soma and neurite imaging by Diffusion MRI. NeuroImage, 215, 116835.

Pfluger, T., Weil, S., Weis, S., Vollmar, C., Heiss, D., Egger, J., Scheck, R., & Hahn, K. (1999). Normative volumetric data of the developing hippocampus in children based on magnetic resonance imaging. Epilepsia, 40(4), 414–423.

Poppenk, J., Evensmoen, H. R., Moscovitch, M., & Nadel, L. (2013). Long-axis specialization of the human hippocampus. Trends in Cognitive Sciences, 17(5), 230–240.

Robillard, K. N., Lee, K. M., Chiu, K. B., & MacLean, A. G. (2016). Glial cell morphological and density changes through the lifespan of rhesus macaques. *Brain*, Behavior, and Immunity, 55, 60–69.

Rossi, R., Arjmand, S., Bærentzen, S. L., Gjedde, A., & Landau, A. M. (2022). Synaptic vesicle glycoprotein 2a: Features and functions. Frontiers in Neuroscience, 16.

Rudrapatna, S., Parker, G., Roberts, J., & Jones, D. Can we correct for interactions between subject motion and gradient-nonlinearity in diffusion MRI. 2018.

Sairanen, V., Leemans, A., & Tax, C. M. W. (2018). Fast and accurate Slicewise outlier detection (SOLID) with informed model estimation for diffusion MRI Data. NeuroImage, 181, 331–346.

Sanides, F. (1970). Functional architecture of motor and sensory cortices in primates in the light of a new concept of neocortex evolution. Advances in Primatology, 1, 137–208.

Sanides, F. (1964). The cyto-myeloarchitecture of the human frontal lobe and its relation to phylogenetic differentiation of the cerebral cortex. Journal Fur Hirnforschung, 6.

Setsompop, K., Kimmlingen, R., Eberlein, E., Witzel, T., Cohen-Adad, J., McNab, J. A., Keil, B., Tisdall, M. D., Hoecht, P., Dietz, P., Cauley, S. F., Tountcheva, V., Matschl, V., Lenz, V. H., Heberlein, K., Potthast, A., Thein, H., Van Horn, J., Toga, A., … Wald, L. L. (2013). Pushing the limits of in vivo diffusion MRI for the Human Connectome Project. NeuroImage, 80, 220–233.

Smith, S. M., Jenkinson, M., Woolrich, M. W., Beckmann, C. F., Behrens, T. E. J., Johansen-Berg, H., Bannister, P. R., De Luca, M., Drobnjak, I., Flitney, D. E., Niazy, R. K., Saunders, J., Vickers, J., Zhang, Y., De Stefano, N., Brady, J. M., & Matthews, P. M. (2004). Advances in functional and structural MR image analysis and implementation as FSL. NeuroImage, 23.

Squire, L. R. (1992). Memory and the hippocampus: A synthesis from findings with rats, monkeys, and humans. Psychological Review, 99(2), 195–231.

Strange, B. A., Witter, M. P., Lein, E. S., & Moser, E. I. (2014). Functional organization of the Hippocampal Longitudinal Axis. Nature Reviews Neuroscience, 15(10), 655–669.

Stüber, C., Morawski, M., Schäfer, A., Labadie, C., Wähnert, M., Leuze, C., Streicher, M., Barapatre, N., Reimann, K., Geyer, S., Spemann, D., & Turner, R. (2014). Myelin and iron concentration in the human brain: A quantitative study of MRI contrast. NeuroImage, 93, 95–106.

Sweatt, J. D. (2010). Hippocampal function in cognition. Mechanisms of Memory, 128–149.

Tamnes, C. K., Bos, M. G. N., van de Kamp, F. C., Peters, S., & Crone, E. A. (2018). Longitudinal development of hippocampal subregions from childhood to adulthood. Developmental Cognitive Neuroscience, 30, 212–222.

Tournier, J.-D., Smith, R., Raffelt, D., Tabbara, R., Dhollander, T., Pietsch, M., Christiaens, D., Jeurissen, B., Yeh, C.-H., & Connelly, A. (2019). MRTRIX3: A fast, flexible and open software framework for medical image processing and visualisation. *NeuroImage*, *202*, 116137. Longitudinal development of hippocampal subregions from childhood to adulthood. Developmental Cognitive Neuroscience, 30, 212–222.

Tustison, N. J., Avants, B. B., Cook, P. A., Zheng, Y., Egan, A., Yushkevich, P. A., & Gee, J. C. (2010). N4ITK: Improved N3 bias correction. IEEE Transactions on Medical Imaging, 29, 1310–1320.

Uchihara, T. (2007). Silver diagnosis in Neuropathology: Principles, practice and revised interpretation. Acta Neuropathologica, 113(5), 483–499.

Uematsu, A., Matsui, M., Tanaka, C., Takahashi, T., Noguchi, K., Suzuki, M., & Nishijo, H. (2012). Developmental trajectories of amygdala and hippocampus from infancy to early adulthood in healthy individuals. PLoS ONE, 7(10).

Veraart, J., Fieremans, E., & Novikov, D. S. (2016). Diffusion MRI noise mapping using random matrix theory. Magnetic Resonance in Medicine, 76(5), 1582–1593.

Vinci-Booher, S., Schlichting, M. L., Preston, A. R., & Pestilli, F. (2023). Development of Human Hippocampal Subfield Microstructure and Relation to Associative Inference. Cerebral Cortex, 33, 10207–10220.

Vos, S. B., Tax, C. M., Luijten, P. R., Ourselin, S., Leemans, A., & Froeling, M. (2016). The importance of correcting for signal drift in diffusion MRI. Magnetic Resonance in Medicine, 77(1), 285–299.

Wierenga, L., Langen, M., Ambrosino, S., van Dijk, S., Oranje, B., & Durston, S. (2014). Typical development of basal ganglia, hippocampus, amygdala and cerebellum from age 7 to 24. NeuroImage, 96, 67–72.

Wolf, D., Fischer, F. U., de Flores, R., Chételat, G., & Fellgiebel, A. (2015). Differential Associations of age with volume and microstructure of hippocampal subfields in healthy older adults. Human Brain Mapping, 36(10), 3819–3831.

Zeineh, M. M., Palomero-Gallagher, N., Axer, M., Gräβel, D., Goubran, M., Wree, A., Woods, R., Amunts, K., & Zilles, K. (2017). Direct visualization and mapping of the spatial course of fiber tracts at microscopic resolution in the human hippocampus. Cerebral Cortex, 27, 1779–1794.

Zhang, H., Schneider, T., Wheeler-Kingshott, C. A., & Alexander, D. C. (2012). NODDI: Practical in vivo neurite orientation dispersion and density imaging of the human brain. NeuroImage, 61(4), 1000–1016.

Zhang, Y., Brady, M., & Smith, S. (2001). Segmentation of brain MR images through a hidden markov random field model and the expectation-maximization algorithm. IEEE Transactions on Medical Imaging, 20(1), 45–57.

