## Supplementary figures for "The developing hippocampus: Microstructural evolution through childhood and adolescence"

### Supplemental Figures for "The developing hippocampus: Microstructural evolution through childhood and adolescence"

[1]Robarts Research Institute, Western University, London, ON, Canada; [2]Centre for Functional and Metabolic Mapping, Western University, London, ON, Canada; [3]Department of Neurosurgery, The Royal Children's Hospital, Melbourne, Australia; [4]Cardiff University Brain Research Imaging Centre (CUBRIC), Cardiff University, Cardiff, United Kingdom; [5]Center for Biomedical Imaging, Department of Radiology, New York University Grossman School of Medicine, New York, NY, United States; [6]School of Computer Science and Informatics, Cardiff University, Cardiff, United Kingdom; [7]Department of Medical Biophysics, Western University, London, ON, Canada

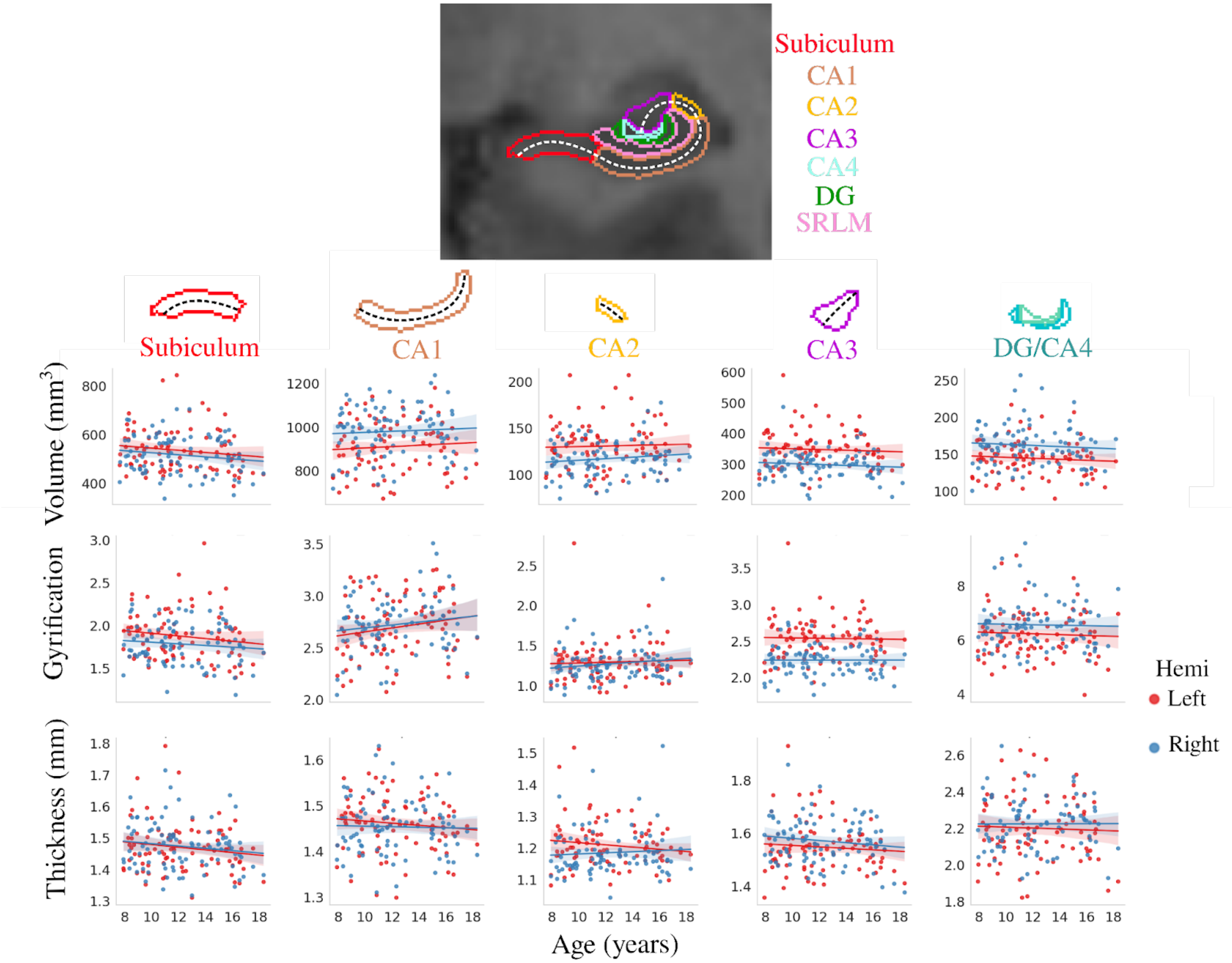


**Supplementary Figure S1.** Relationship between age and subfield averaged macrostructural measures of volume, gyrification, and thickness between the left and right hemisphere. The dashed lines approximately represent the midthickness surface which gyrification and thickness were calculated and then averaged on. CA - cornu ammonis; DG - dentate gyrus; SRLM - stratum radiatum lacunosum moleculare.

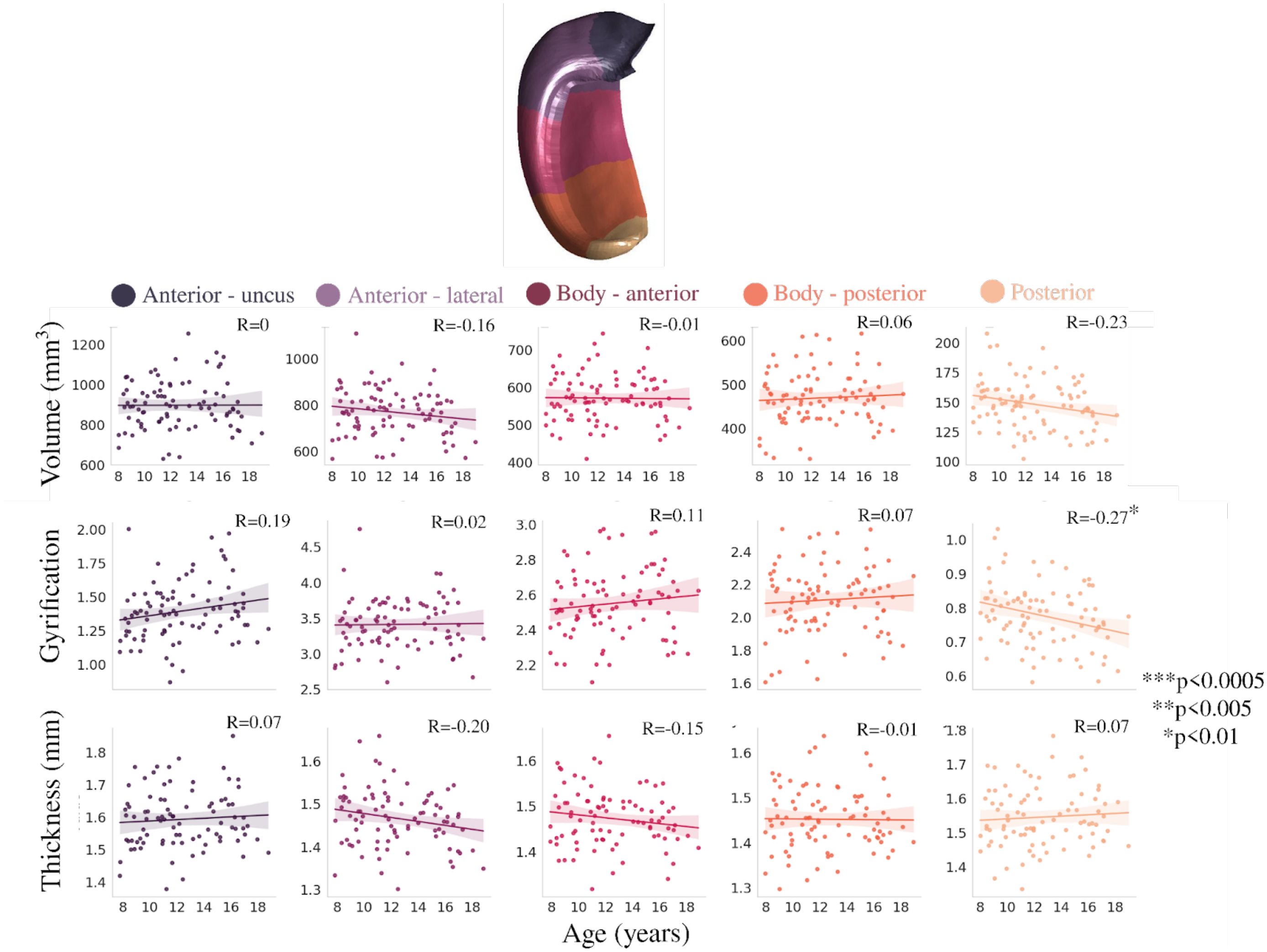


**Supplementary Figure S2.** Relationship between age and long-axis averaged macrostructural measures of volume, gyrification, and thickness.

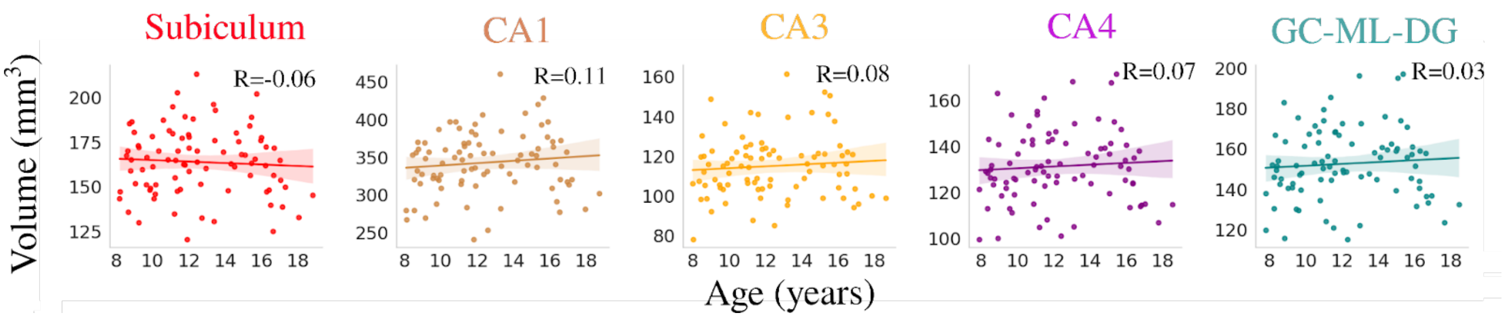


**Supplementary Figure S3.** Relationship between age and subfield volume derived from FreeSurfer.

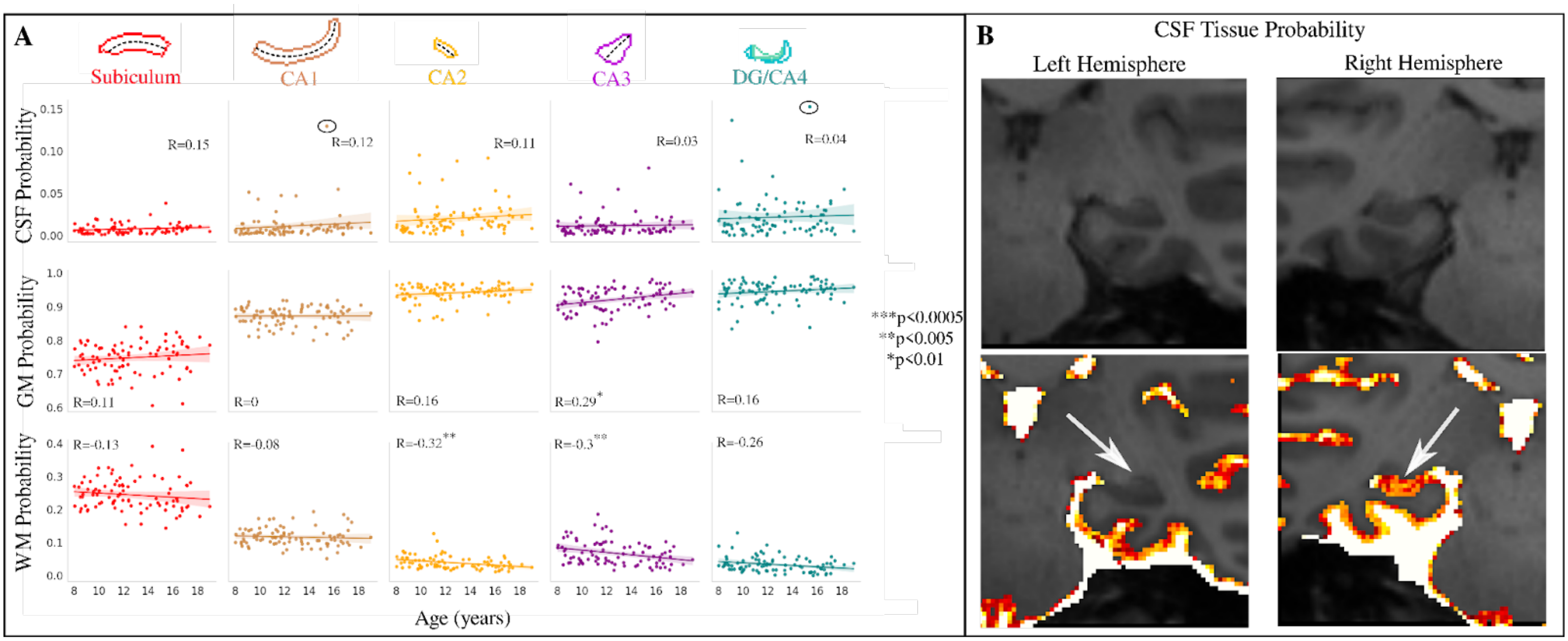


**Supplementary Figure S4.** (A) Relationship between age and subfield averaged partial volume measures of CSF, GM, and WM. The circled points represent a subject with a high probability of CSF. (B) T1w image and CSF probability output for the circled subject in (A). The right hemisphere has a misestimated high CSF probability. CA - cornu ammonis; DG - dentate gyrus; SRLM - stratum radiatum lacunosum moleculare.

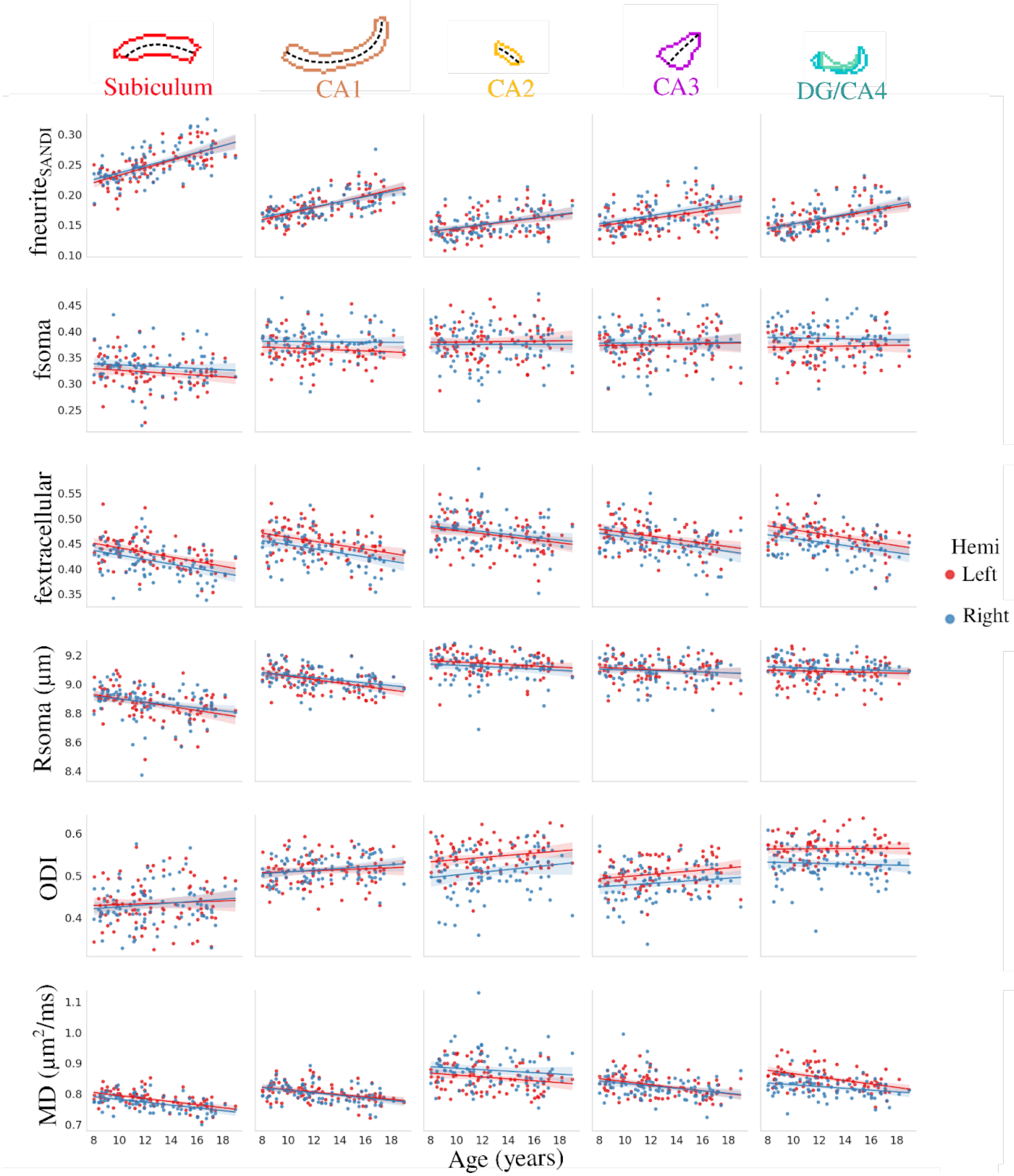


**Supplementary Figure S5.** Relationship between age and subfield averaged microstructural measures of neurite ($fneurite_{SANDI}$), soma (fsoma), and extracellular (fextracellular) signal fractions, soma radius, orientation dispersion index (ODI) and mean diffusivity (MD) between the left and right hemisphere. The dashed lines approximately represent the midthickness surface which the metrics were sampled and then averaged on. CA - cornu ammonis; DG - dentate gyrus.

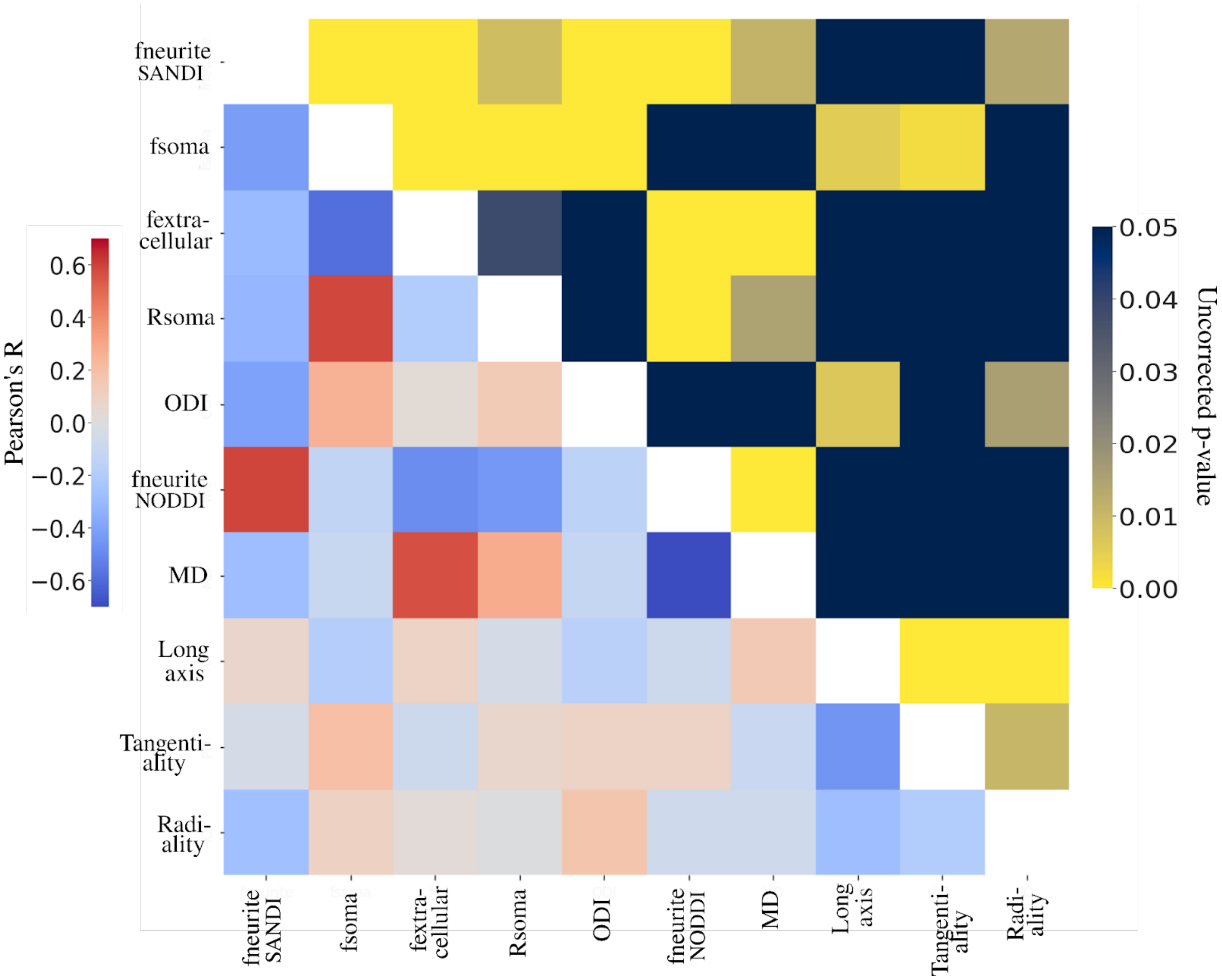


**Supplementary Figure S6.** Spin test correlation between all the age contrasted t-statistic microstructure maps which capture vertex-wise age-related microstructural changes.

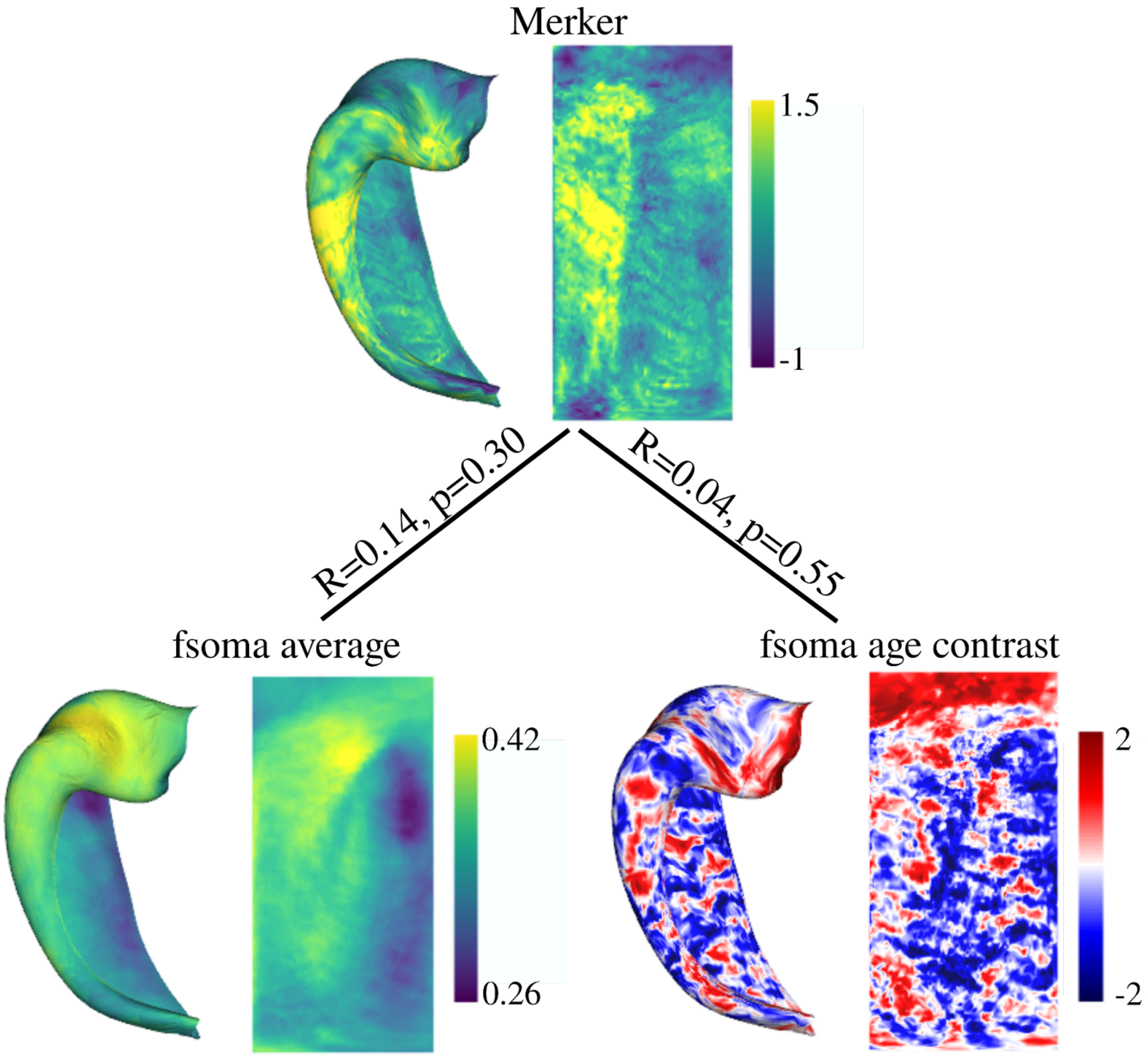


**Supplementary Figure S7.** Correlating a Merker stain for cell bodies with fsoma derived from SANDI (DeKraker et al., 2024; Amunts et al., 2013). Bottom left is fsoma averaged across all subjects (i.e. averaging across age) and bottom right is the age contrasted t-statistic map capturing vertex-wise age-related changes of fsoma.

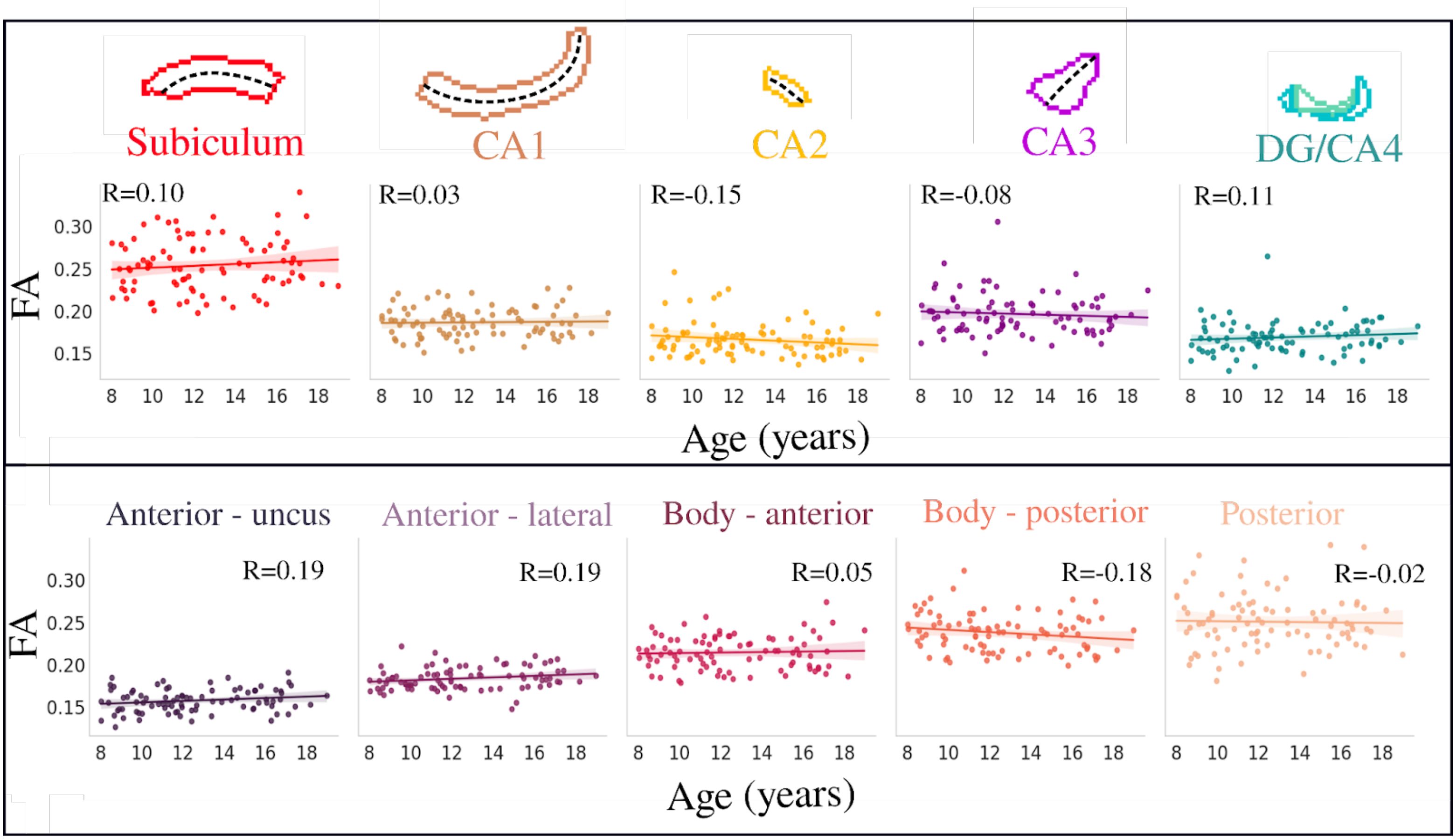


**Supplementary Figure S8**. Correlating age with subfield (top) and long-axis (bottom) averaged fractional anisotropy (FA) derived from DTI.
